# Long-Term Ecological Responses of a Dipterocarp Forest to Climate Changes and Nutrient Availability

**DOI:** 10.1101/2022.12.04.518039

**Authors:** Ana Prohaska, Alistair W.R. Seddon, Oliver Rach, Andrew Smith, Dirk Sachse, Katherine J. Willis

## Abstract

- Understanding the long-term impact of projected climate change on tropical rainforests is critical given their central role in the Earth’s system. Palaeoecological records can provide a valuable perspective on this problem. Here we examine for the first time the effects of past climatic changes on the dominant forest type of Southeast Asia – Dipterocarp forest.
- We use a range of proxies extracted from a 1,400-year-old lacustrine sedimentary sequence from north-eastern Philippines to model long-term vegetation responses of Dipterocarp forest, including its dominant tree group Dipterocarps, to changes in precipitation, fire and nutrient availability over time.
- Our results show a positive relationship between Dipterocarps pollen accumulation rates (PARs) and leaf wax hydrogen isotope values, which suggests a negative effect of drier conditions on Dipterocarp tree abundance. Furthermore, we find a positive relationship between Dipterocarp PARs and the proxy for phosphorus availability, which suggests phosphorus controls the productivity of these keystone trees on longer time scales. Other plant taxa show widely varying relationships with the abiotic factors, demonstrating a high diversity of functional responses.
- Our findings provide novel insights into Dipterocarp forest responses to changing climatic conditions in the past, and highlight potential impacts of future climate change on this globally important ecosystem.

## Introduction

Lowland tropical rainforests make up the most productive terrestrial biome and cover nearly 10% of Earth’s land surface (Saatchi *et al*. 2011). As such, they represent a key component of the models predicting Earth’s capacity to combat increasing levels of CO_2_ (Bonan & Levis 2010; Huntingford *et al*. 2013). Yet the nature and extent to which climate and nutrient availability influence the productivity and reproduction of tropical rainforests remain poorly resolved (Cleveland *et al*. 2011). This is particularly true for the Dipterocarp forests that once covered much of peninsular Malaysia, Sumatra, Java, Borneo, and the Philippines (Appanah 1985). Due to record high rates of deforestation, they presently occupy only 30% of their original range (Sodhi *et al*. 2004; Wilcove *et al*. 2013). Dipterocarp forests are ecologically unique among tropical rainforests in that they engage in interannual mass flowering, and are overwhelmingly dominated by a single tree family, Dipterocarpaceae, whose members constitute up to 10% of total tree diversity, over 50% of the basal area and nearly 80% of all emergent trees in these forests (Appanah 1985).

Observational studies have shown that drought and fire events can exert a large influence on the ecological processes in Dipterocarp forests on annual scales. Severe drought events in the region, of which El Niño-Southern Oscillation (ENSO) is the main driver, lead to increased mortality for most tree species in Dipterocarp forests, with large trees being particularly susceptible. For instance, in a single drought event in 1998, large trees (i.e., >80 cm d.b.h.) made up more than 45% of total stem mortality in Dipterocarp forests (Van Nieuwstadt *et a*l. 2005). Furthermore, fire events, which are more likely to occur during severe droughts, can cause near total mortality of smaller trees (Van Nieuwstadt *et a*l. 2005). The resulting increase in tree mortality can lead to higher occurrence of forest gaps, which presently form only a small portion of the total area (Ghazoul and Sheil 2010). It has therefore been hypothesized that long-term changes in the magnitude and frequency of drought and fire events, associated with the changes in ENSO activity projected for this century, will profoundly alter the structure of Dipterocarp forests (Whitmore 1988; Ghazoul 2016).

In contrast, other work has suggested that El Niño-induced drought events play a key role in the reproduction of Dipterocarp forests because they trigger mass flowering events (Shoko Sakai *et al*. 2006). Most Dipterocarp species reproduce only during such events. For example, more than 60% of *c*. 200 Dipterocarp species in North Borneo were observed to flower during a single strong El Niño drought event in 1955 (Woods 1956), and almost 50% of the mature Dipterocarp individuals have been shown to flower during mass flowering years (Burgess 1972; Cockburn 1975). Mass flowering of Dipterocarps are often accompanied by heavy flowering of many other plant species, including members of Anacardiaceae, Euphorbiaceae, Annonaceae, Fabaceae, Burseraceae, Moraceae and Myristicaceae (Appanah 1985). An alternative hypothesis, therefore, is that rather than a decline in Dipterocarp tree abundance, stronger El Niño-related drought regimes predicted for this century will instead lead to stronger/ more frequent flowering events, boosting the reproduction of Dipterocarp forests.

In addition to precipitation and fire, understanding nutrient limitations in Dipterocarp forests is essential for predicting their responses to current and future disruptions in nutrient cycling (Sala *et al*. 2000; Boisvenue & Running 2006; Hou *et al*. 2018). Phosphorus is a primary plant macronutrient with particularly low concentrations in tropical soils and it is thus widely assumed to be the main limiting nutrient to the productivity of lowland tropical rainforests (Vitousek 1984) (in contrast to nitrogen that is believed to be generally abundant due to biological N fixation (Jenny 1950; Cleveland *et al*. 1999; Galloway *et al*. 2004). However, evidence of phosphorus limitation is scarce. Previous work has indicated that the response of tropical trees to increased phosphorus supply is highly heterogeneous, depending upon light conditions (Thompson *et al*. 1988), successional status (Lawrence 2003), mycorrhizal association (Moyersoen *et al*. 1998) and size class (Wright *et al*. 2011; Alvarez-Clare *et al*. 2013). Previous studies also found a weak relationship between primary productivity and phosphorus availability (Cleveland *et al*. 2011), with nitrogen addition eliciting a tree growth response as large as phosphorus addition (Wright *et al*. 2018). These findings come predominantly from short-term experimental studies at stand-level (Vitousek *et al*. 1993; Mirmanto *et al*. 1999; Newbery *et al*. 2002; Davidson *et al*. 2004; Wright *et al*. 2011; Alvarez-Clare *et al*. 2013; Cunha *et al*. 2022) and at the level of individual trees (Burslem *et al*. 1996; Gunatilleke *et al*. 1997; Lawrence 2003; Yavitt & Wright 2008), and from observational studies across environmental gradients that vary in soil nutrient availability (Vitousek *et al*. 1992; Santiago *et al*. 2004; Santiago & Mulkey 2005; Baltzer *et al*. 2008; Fyllas *et al*. 2009). However, a large portion of the biomass production in a tropical rainforest stand is carried out by large trees (Rutishauser *et al*. 2010; Slik *et al*. 2010; Ferry Slik *et al*. 2013), and according to a hypothesis by Vitousek (1984), it is these individuals that are most limited by phosphorus availability. Thus, understanding the response of a tropical rainforest to phosphorus availability is not possible without understanding the response of its large trees. Given the long life span (multi-centennial) and generation time (multi-decadal) of large rainforest trees, predicting future responses of tropical rainforests to changing conditions is extremely challenging. However, distinct changes in precipitation amount, some of which linked to ENSO dynamics, along with changing fire intensity and nutrient availability, have been recorded over the past millennia (Cobb *et al*. 2003; Conroy *et al*. 2008; Overpeck & Cole 2010; Cobb *et al*. 2013; McLauchlan *et al*. 2013; Marlon *et al*. 2013; Prohaska *et al*. 2022). Comparing paleoclimate records with fossil pollen and other fossil proxies that capture plant responses to these changes therefore holds great potential for understanding and predicting the effects of future climate change on Dipterocarp forests.

Here, we investigate the long-term vegetation dynamics of Dipterocarp forests from a 1,400-year-old laminated sedimentary sequence from Bulusan Lake in the north-eastern Luzon, Philippines (Fig. 1a-c). To that end, we generate bi-decadal pollen accumulation rates (PARs, grains cm^−2^yr^−1^) (Davis & Deevey 1964; Giesecke & Fontana 2008) and use the PARs as an independent proxy of plant taxa abundance (see Supporting Information). We also generate a record of carbon isotope values of leaf wax alkanes (δ^13^C_wax_, ‰), and use *n*C_29_ alkane δ^13^C values (hereafter δ^13^C_wax_ record) as an independent proxy of the ratio of C3 vegetation (trees and grasses) to C4 vegetation (grasses). We examine the response of individual Dipterocarp forest plant taxa, including the dominant Dipterocarp trees, and the overall forest vegetation cover, to a range of climatic and nutrient factors. A recently generated record of hydrogen isotope values of leaf wax alkanes (δD_wax_ ‰), from the Bulusan sedimentary sequence has demonstrated high precipitation variability throughout the last 1,400 years, as well as a prolonged period of El Niño-like conditions during the Little Ice Age (1630-1900 AD) (Prohaska *et al*. 2022). Hence, the Bulusan sequence presents an excellent opportunity to examine the effects of changing precipitation conditions on Dipterocarp forests. We use *n*C_29_ alkane δD values from the Bulusan sequence (hereafter δD_wax_ record) as a proxy record of precipitation variability related to changes in ENSO mean state conditions (Prohaska *et al*. 2022). We also generate fossil macrocharcoal influx rates (particles cm^−2^yr^−1^) as a proxy record of local fire activity (Clark 1988; Mooney & Tinner 2010), and stable nitrogen isotope composition of bulk sediments (sedδ^15^N, ‰) and phosphorus concentration of bulk sediments (sedP_conc_, XRF counts per second) (Croudace *et al*. 2006) as proxy records of nitrogen and phosphorus availability, respectively. Using Generalised Additive Models (GAMs), we first examine the temporal trends in PARs, particularly in relation to the prolonged period of El Niño-like conditions. Following this, we employ GAMs to evaluate the relationship between PARs and abiotic factors on a bi-decadal scale.

**Fig. 1.**
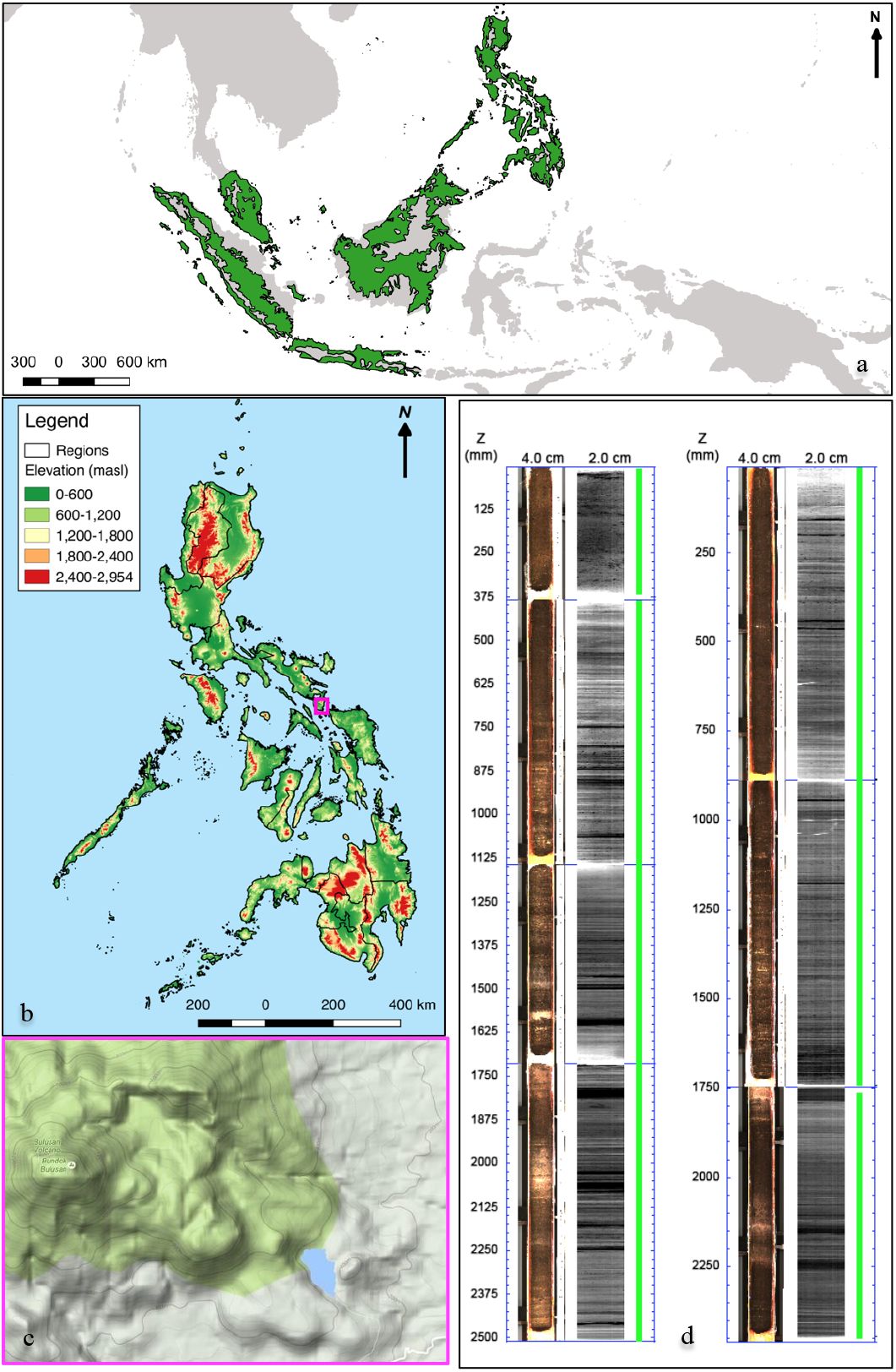
Geographical location and sedimentary record from the Bulusan site. **a,** Map showing the geographic distribution of Dipterocarp forests (in green). Data from Olson *et al*. (2001), map generated in QGIS version 2.14.3 (QGIS Development Team 2016). **b,** Topographic map of the Philippines showing the location of the study site. Data from PhilGIS (2016), map generated in QGIS version 2.14.3. **c,** Map showing Bulusan Lake (in blue) within the larger landscape, with Mt Bulusan volcano in the west (in green) (Google Maps 2016). **d,** Optical and radiographic profiles of the sedimentary record obtained from the Bulusan Lake. The radiograph images depict the level of organic content, with higher values shown as lighter bands. The green line represents the validation of the measurement quality along the cores. Images were generated in the ItraxPlot program (Croudace & Rothwell 2015).

## Methods

### Study site

Bulusan Lake is a 0.28 km^2^ lake lying at 360 m.a.s.l. at the foot-slopes of Mt Bulusan, a stratovolcano at the southeast end of the Bicol Volcanic Arc (Delfin *et al*. 1993). The lake consists of two basins: a 12 m deep round basin, and a larger, 24 m deep basin of more narrow profile. The two basins have no inlets or outlets and comprise a closed system fed primarily by precipitation and groundwater. The waters of Bulusan Lake are of clear, bright green colour and no floating or submerged vegetation has been recorded at the lake. The lake is part of the Bulusan Volcano National Park, a nature reserve that was first established in 1935, and where logging and hunting activities are strictly regulated (Abada Gatumbato *et al*. 2011).

The region has a dominant rainfall season occuring during autumn, and is a member of global “Autumn Monsoon precipitation” region (Ramesh *et al*. 2021). The western North Pacific monsoon also operates over the region, however its contribution to the total annual rainfall is smaller. In addition, typhoons are a source of episodic precipitation, accounting for up to 30% of northeastern Philippines rainfall in a year (Rodgers *et al*. 2000). ENSO is the main driver of inter-annual rainfall variability at Bulusan, resulting in reduced (increased) precipitation at Bulusan during El Niño (La Niña) years. This makes the Bulusan record ideally suited to provide insights into the vegetation responses to mean changes in ENSO dynamics on longer time scales.

The vegetation surrounding Bulusan Lake is classified as Lauan Dipterocarp forest dominated by Dipterocarp tree *Shorea contorta* (Abada Gatumbato *et al*. 2011). In addition to *Shorea*, other notable large tree genera in the Bulusan Dipterocarp forest include Dipterocarps trees *Vatica* and *Parashorea*, as well as *Pterocarpus, Dracontomelon, Agathis, Pinanga* and *Ficus* (Abada Gatumbato *et al*. 2011). Species such as *Litsea, Aglaia, Artocarpus*, and *Syzygium* dominate in the lower canopy, while *Elaeocarpus, Leea* and *Bischofia* are prominent subcanopy trees. Presently, pioneer plants comprise only a small fraction of the Dipterocarp forest vegetation because their successful establishment is limited to gaps (Ashton & Seidler 2014) which cover only around 1% of total forest area (Ghazoul 2016). Common pioneer taxa include *Macaranga, Trema, Melastoma, Dillenia, Alstonia*, Poaceae and Polypodiopsida (Woods 1989; Ashton & Seidler 2014; Davies & Ashton 1999; Co *et al*. 2006).

### Collection of lake sediments

Two overlapping sedimentary sequences, BUL1 and BUL2, were collected as 1 m cores in March 2013 from the smaller basin of Bulusan Lake with a Livingstone Piston corer from an anchored platform. The sediments were shipped to the University of Oxford and stored at 5°C. The two sedimentary sequences were used to create a continuous, undisturbed composite sequence using (non-annual) laminations present throughout the sequence (Fig. 1d). Sediment laminations were mapped using optical and radiograph images taken with an Itrax Core Scanner at the University of Aarhus in September 2013 (Fig. 1d). For more information about the Bulusan laminations, please see Prohaska *et al*. (2022). All proxy records in this study come from these two sedimentary sequences and are constrained by a high-resolution age-depth model (see Supporting Information for more details).

### Extraction and processing of fossil proxies

Sediment samples were extracted from the top 3.0 m of the composite Bulusan sequence as a continuous series of 4 cm long slices of 1 cm^3^ volume, with each slice corresponding to a ~20-year interval. Standard protocols were employed for the extraction and counting of fossil pollen (Bennett and Willis 2001), and a reference collection of pollen types from the region was generated using herbarium samples, and combined with printed and online sources (e.g., Australasian Pollen and Spore Atlas Online Database (APSA Members 2007), Pollen flora of the Philippines (Jagudilla-Bulalacao 1997), Pollen Flora of Taiwan (Huang 1972)). Five hundred terrestrial pollen grains were counted and identified in each sample. PARs were calculated as the number of pollen grains deposited in 1 cm^2^ of sediment per year according to the standard procedures (Stockmarr 1971) and used to infer changes in the abundance of plant taxa over time. For obtaining macrocharcoal influx rates as a proxy of local fire activity, we calculated the total number of macrocharcoal particles (>150 μm) identified in sediment samples under a Nikon stereoscope at 18x magnification (Clark 1988; Mooney & Tinner 2010).

### n-alkane paleohydrological and paleovegetation records

For detailed description of extraction, identification and quantification of higher leaf wax *n*-alkanes (*n*C_23-33_ homologues) and the compound-specific hydrogen isotope measurement and evaluation, please see Prohaska *et al*. (2022).

Compound-specific stable carbon isotope ratios of the aliphatic fraction were measured at the German Research Centre for Geosciences in Potsdam using a ThermoFisher Delta-V-Plus Isotope Ratio Mass Spectrometer coupled to a ThermoFisher TraceGC 1310 gas chromatograph. The samples were measured on a 30 m DB5-MS column with an inner diameter of 0.25 mm and a film thickness of 0.25 μm. The samples were measured with the following GC-temperature program: i) temperature was held for 2 mins at 70 °C, ii) temperature was increased to 150 °C at a rate of 15 °C per min, iii) oven temperature was increased to 320 °C at 5 °C per min, and iv) the final temperature was held for 10 mins. A standard containing *n*C_16_ to *n*C_30_ alkanes with known δD values (mix A6, Arndt Schimmelmann, University of Indiana) was measured in duplicate at the beginning and the end of each sequence or after six sample injections, and then used for normalisation of δD values to the Vienna Standard Mean Ocean Water (VSMOW) scale. The alkane standard mix was measured with the same temperature program as the samples. Samples and standards were measured in duplicates. The mean standard deviation of all measured samples for the *n*C_23_, *n*C_25_, *n*C_27_, *n*C_29_, *n*C_31_ and *n*C_33_ alkanes (n=41 to 65) was approximately 0.3‰, 0.3‰, 0.2‰, 0.1‰, 0.1‰ and 0.3‰, respectively.

### Bulk sediment nitrogen isotope composition

Bulk sediment samples from the Bulusan cores were analysed for δ^15^N, along with percentage carbon, percentage nitrogen, and δ^13^C, at the Godwin laboratory for Palaeoclimate Research, Department of Earth Sciences, University of Cambridge. The dried and homogenised samples were carefully weighed into a tin capsule (c. 10 mg), sealed, and loaded into the autosampler. The samples were analysed using a Costech Elemental Analyser attached to a Thermo DELTA V mass spectrometer in continuous flow mode. Weighted samples of standards were analysed at various points throughout the run allowing percentage nitrogen and percentage carbon to be calculated for the batch of samples. Reference standards from IAEA in Vienna were also run at intervals throughout the sequence and these values were used to calibrate to the international standards for ^14^N/^15^N (δ^15^N air) and ^12^C/^13^C (δ^13^C VPDB). Precision of analyses is +/- 0.5% for C and N, and better than 0.1 permille for δ^13^C and δ^15^N.

### Bulk sediment phosphorus concentration

We conducted high-resolution analysis of major and trace chemical elements, including phosphorus, in the Bulusan sedimentary sequence using the ITRAX Core Scanner (Croudace *et al*. 2006) at Aarhus University, Denmark. The cores were scanned at a step-size of 0.2 mm to obtain an ED-XRF spectrum at each step. A digital continuous-strip X-radiograph, 22 mm in width, was collected immediately prior to the XRF scan using a charge-coupled line camera. The count time for each measurement was 30 seconds to ensure that minor elements are well detected. The X-ray beam used to irradiate the cores is generated using a 3kW Mo X-ray tube run at 55kV and 50 ma for x-radiography and 30kV and 50ma for the XRF scan. The recorded spectra were deconvolved in live time to construct profiles of peak area integrals for individual elements. The element peak area reflects element abundances within the sediment. For more details on the analysis and interpretation of sedP_conc_, please see Supporting Information.

### Generalised additive modelling

The temporal trends in mean and variance of the generated PARs records were estimated using Generalised Additive Models (GAMs) following (Simpson 2018). GAMs are statistical models that can be used to estimate trends in mean and variance as smooth functions of time using automatic smoothness selection methods that objectively determine the complexity of the fitted trend (Hastie & Tibshirani 1986; Hastie & Tibshirani 1990; Wood 2017). GAMs are particularly powerful for analysing palaeoenvironmental time series because they allow to estimate complex, non-linear trends and to identify time intervals of significant change. Two GAM models were applied in R version 3.6.3 (R Core Team 2020) using the mgcv 1.8-31 package (Wood 2018): gam, that accounts for heteroscedasticity, and gamm+car(1), that accounts for the car(1) correlation. In both GAM models we used tweedie distribution to limit predicted values to positive numbers. We checked if the size of the basis expansion is sufficient by fitting a different number of basis functions (k) using gam.check function. For the gam models, we tested and, where appropriate, accounted for heteroscedasticity that may have arisen due to variation in sedimentation rates over time, and thereby, in the span of time that each sample represents. The fitted GAM smoother was considered significant at p-value <0.05. Both gamm+car(1) and gam models for individual PAR datasets were plotted with confidence intervals that were generated using the critical value from the t-distribution.

We also used GAMs to assess the effect of abiotic factors (δD_wax_, macrocharcoal influx rates, sedimentary δ^15^N and P_conc_) on the plant taxa abundance (PARs) and C3/ C4 vegetation composition (δ^13^C_wax_). Macrocharcoal influx rates were transformed using square root function to reduce effect of a small number of disproportionately large values (log function not used due to many zero values). Two GAMs were applied as described above. In both GAMs for PARs we used a Gamma distribution to limit predicted PAR values to positive numbers (tweedie family was also attempted but the models did not converge over similar ρ values). Since the Gamma family does not support zero values, these were removed from PAR datasets prior to the analysis. In the macrocharcoal influx rates and P_conc_ datasets, two points identified by preliminary analysis as strong outliers were removed prior to modelling. For the δ^13^C_wax_ record, we used a Gaussian distribution. The two models (gam and gamm+car(1)) both included all four abiotic factors as explanatory variables, and were compared according to their Akaike Information Criterion (AIC) coefficients. The best model was selected based on the lowest AIC coefficients, with a simpler model selected whenever the difference in AIC coefficients between two models was smaller than 2. Explanatory variables were considered significant at the 95% level of confidence. To evaluate the impact of removing zero values from the PAR records, we also run GAM based on the Quasi-Poisson family that showed similar results (see Supporting Information). The Quasi-Poisson family is suitable for overdispersed count data and supports the presence of zero values in data, but requires the conversion of PAR values to integers. Where a model with log link for Gamma and/ or Quasi-Poisson distribution could not converge, square-root (sqrt hereafter) or identity links were used instead, and the converging model with the lowest AIC value was selected.

## Results & Discussion

### Response of Dipterocarp forest vegetation to long-term environmental changes

The δD_wax_ record covering the past 1,400 years indicates that the Bulusan region has experienced a prolonged period of El Niño-like conditions during the late Little Ice Age, i.e., 1630-1900 AD, resulting in a reduction in precipitation evident from an abrupt increase in δD_wax_ values during this time (Prohaska *et al*. 2022) (Fig. 2a). Furthermore, our new record of fossil macrocharcoal influx rates from Bulusan site shows that this period was accompanied by increased fire activity in comparison to the more recent times, though there was a period of even higher fire activity preceding it (1400-1600 AD) (Fig. 2b). Nitrogen isotopic values show limited variability over the last 1,400 years, apart from a pronounced increase at ~1000 AD, followed by a modest decrease to present-day values at ~1250 (Fig. 2c). In contrast, the record of P_conc_ varied extensively at Bulusan over the same period, with notable peaks at ~900 AD and ~1700 AD, and troughs at ~750 AD and ~1200 AD (Fig. 2d).

**Fig. 2.**
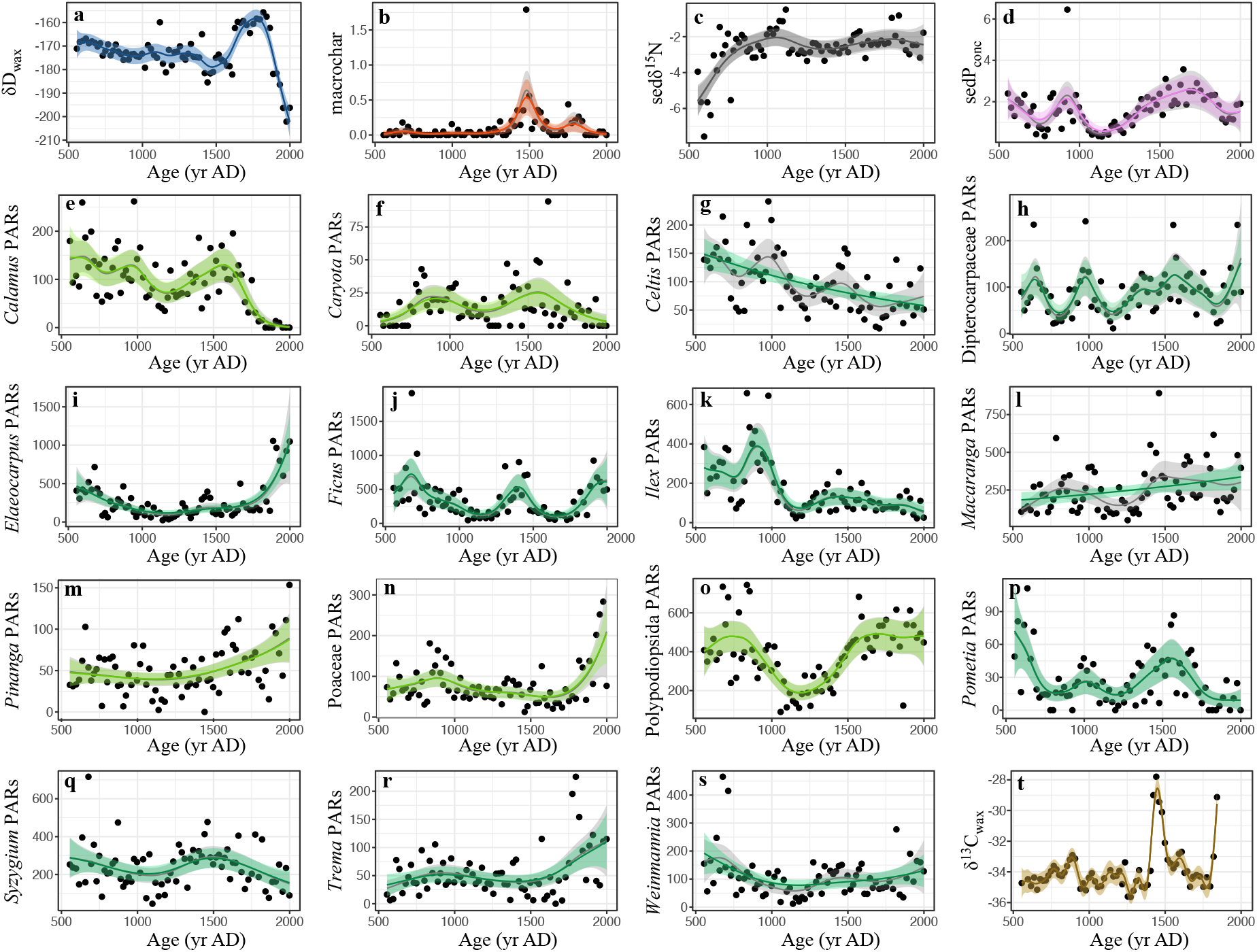
The temporal trends in mean and variance for the key paleo-records from Bulusan Lake according to the best fitted generalised additive models. For the two models shown, the colour fit is the result of a GAM with a continuous-time AR(1) process estimated using REML smoothness selection (gamm+car(1) = ~ Age), while the grey fit is that of a simple GAM with GCV-based smoothness selection (gam = ~ Age)). Light green depicts predominately pioneer taxa, and dark green depicts predominately shadow-tolerant taxa. The fitted GAM smoother is overall significant (p-value <0.05). PARs = pollen accumulation rates (grains cm^−2^yr^−1^), macrochar = macrocharcoal influx rates (particles m^−2^yr^−1^), δ^13^C_wax_ = δ^13^C values of the *n*C_29_ alkane (‰), δD_wax_ = δD values of the *n*C_29_ alkane (‰), sedδ^15^N = δ^15^N values of bulk sediment (‰), sedP_conc_ = phosphorus concentration of bulk sediment (XRF counts per second), Age = age of the samples measured as calibrated years AD. Data was analysed using the mgcv 1.8-31 package (Wood 2018) and plotted with *ggplot2* package (Wickham 2009) in R version 3.6.3 (R Core Team 2020).

We find that Dipterocarp PARs exhibited relatively large fluctuations throughout the Bulusan record (Fig. 2h). Notably, Dipterocarp PARs declined and remained low during the 1630-1900 AD period of reduced precipitation. However, the PARs also exhibited at least two comparable declines in abundance prior to this, specifically during 1100-1300 AD and 700-900 AD, despite relatively stable precipitation during these time periods. Furthermore, during the period of most intense fire intensity recorded in the Bulusan record, 1400-1600 AD, Dipterocarp PARs were at some of their highest levels. Finally, the peaks and troughs of Dipterocarpaceae PARs correspond closely to the peaks and troughs in P_conc_. As a result, it is difficult to discern based on the analysis of temporal trends alone the relative contribution of precipitation, fire and nutrient variability to the dynamics of Dipterocarp PARs at Bulusan over the last 1,400 years.

Other plant taxa show widely divergent PAR trends over the last 1,400 years (Fig. 2), and many of the PAR records include significant periods of change according to the first derivatives analysis (Fig. S7). For instance, *Calamus, Pometia* and *Syzygium* show a decrease in PAR values during the 1630-1900 AD period while *Pinanga* and *Trema* show the opposite. Furthermore, *Macaranga, Pometia*, and *Syzygium* show a distinct increase in PAR values during the peak in charcoal values between 1400-1600 AD, while the PAR values of *Calamus, Caryota*, and *Pometia* closely follow the P_conc_ changes over time. One common theme among several PARs is an increase to present day. For instance, the post-1800 AD decrease in PARs of *Calamus* and *Caryota* that may be at least partially linked to anthropogenic activities, i.e., harvesting for materials and food, that have been documented in this area (see Supporting Information). Moreover, a pronounced post-1900 AD increase in PARs of *Elaeocarpus, Trema* and Poaceae coincides with the onset of more intense anthropogenic activities in the area, including moderate logging around the Bulusan Lake (Abada Gatumbato *et al*. 2011). While land use change leading to a more open environment is a possible explanation to the recent PAR changes in these predominantly pioneer taxa, it is not included directly in our analysis due to lack of independent records for these activities. Interestingly, these activities appear not to have had a notable effect on Dipterocarp abundance, as their PARs continue an upward trajectory. Overall, our results reveal a high diversity of responses across different plant taxa, which is to be expected for this forest type.

Values of δ^13^C_wax_ in the Bulusan record range from −35.6‰ to −27.8‰. Previous studies established that δ^13^C_wax_ values of higher plant n-alkanes reflect their carbon fixation pathways and can thus be used to track changes in C3/ C4 composition of vegetation (e.g., Schefuss *et al*. (2003), Castañeda *et al*. (2009), Tjallingii *et al*. (2011), Garcin *et al*. (2014)). To that end, n-C_29_ and n-C_31_ alkanes have been most widely used because they are the most abundant and ubiquitous n-alkane homologues in both plant material and sedimentary archives. In tropical settings, C3 (trees and shrubs) and C4 (grasses) vegetation produces waxes with average δ13C_wax_ values of −35‰ and −18‰, respectively (Garcin *et al*. 2014). Bulusan δ^13^C_wax_ values are well within the range of C3 vegetation, providing further support that forest cover dominated the surrounding landscape throughout the last 1,400 years with a varying, but overall small, C4 vegetation component. Nevertheless, the Bulusan δ^13^C_wax_ record also shows a marked and abrupt rise in values close to 1500 AD, which closely coincides with the peak in the fossil macrocharcoal influx rates values. This suggests that the enhanced fire activity around this time has led to an increase in the proportion of C4 vegetation to C3 vegetation, perhaps by creating a more open habitat that was suitable for grasses.

For a more detailed description of the temporal trends analysis, please see Supporting Information (Tables S3-S4).

### Influence of climate and nutrient availability on Dipterocarp trees

Our results suggest that on a bi-decadal scale the changes in Dipterocarp PARs are associated with both precipitation and nutrient availability. Specifically, the best-fitted GAM model yielded a significant positive effect (p-value >0.05) of the δD_wax_ values and P_conc_ values on Dipterocarp PARs (Table 1 and Fig. 3, for a more detailed description of the GAM results, please see Supporting Information). This suggests that on multidecadal scales the Dipterocarp abundance is inversely associated with ENSO-mediated drought but positively associated with phosphorus availability.

**Table 1.**
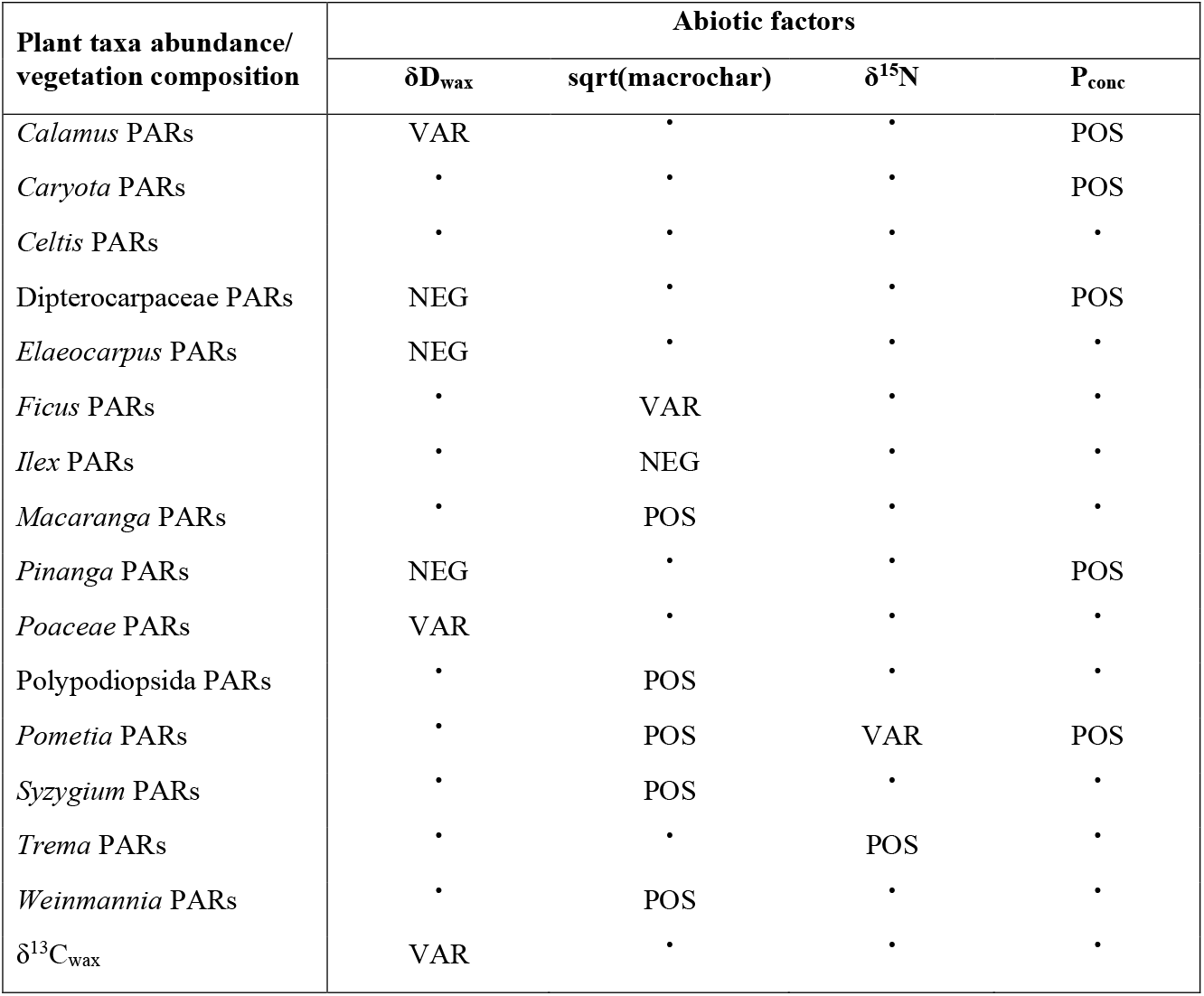
Summary of results of the best fitted generalised additive models for describing the effects of abiotic factors on plant taxa abundance (pollen accumulation rates, PARs) and C3/ C4 vegetation composition ((δ^13^C values of the *n*C_29_ alkane, δ^13^C_wax_) at Bulusan. General model formula: PAR ~ s(δD_wax_) + s(sqrt(macrochar)) + s(sedδ^15^N) + s(sedP_conc_), distribution family = Gamma, smoothing parameter estimation method = Restricted Maximum Likelihood; and δ^13^C_wax_ ~ s(δD_wax_) + s(sqrt(macrochar)) + s(sedδ^15^N) + s(sedP_conc_), distribution family = Gaussian, smoothing parameter estimation method = Restricted Maximum Likelihood. PARs = pollen accumulation rates (grains cm^−2^yr^−1^), macro = macrocharcoal influx rates (particles m^−2^yr^−1^), δ^13^C_wax_ = δ^13^C values of the *n*C_29_ alkane (‰), δD_wax_ = δD values of the *n*C_29_ alkane (‰), sedδ^15^N = δ^15^N values of bulk sediment (‰), sedP_conc_ = phosphorus concentration of bulk sediment (XRF counts per second), POS = significant positive relationship, NEG = significant negative relationship, VAR = significant varying relationship, • = no significant relationship. The fit is considered significant at p-value <0.05. Data was analysed using the mgcv 1.8-31 package (Wood 2018).

**Fig. 3.**
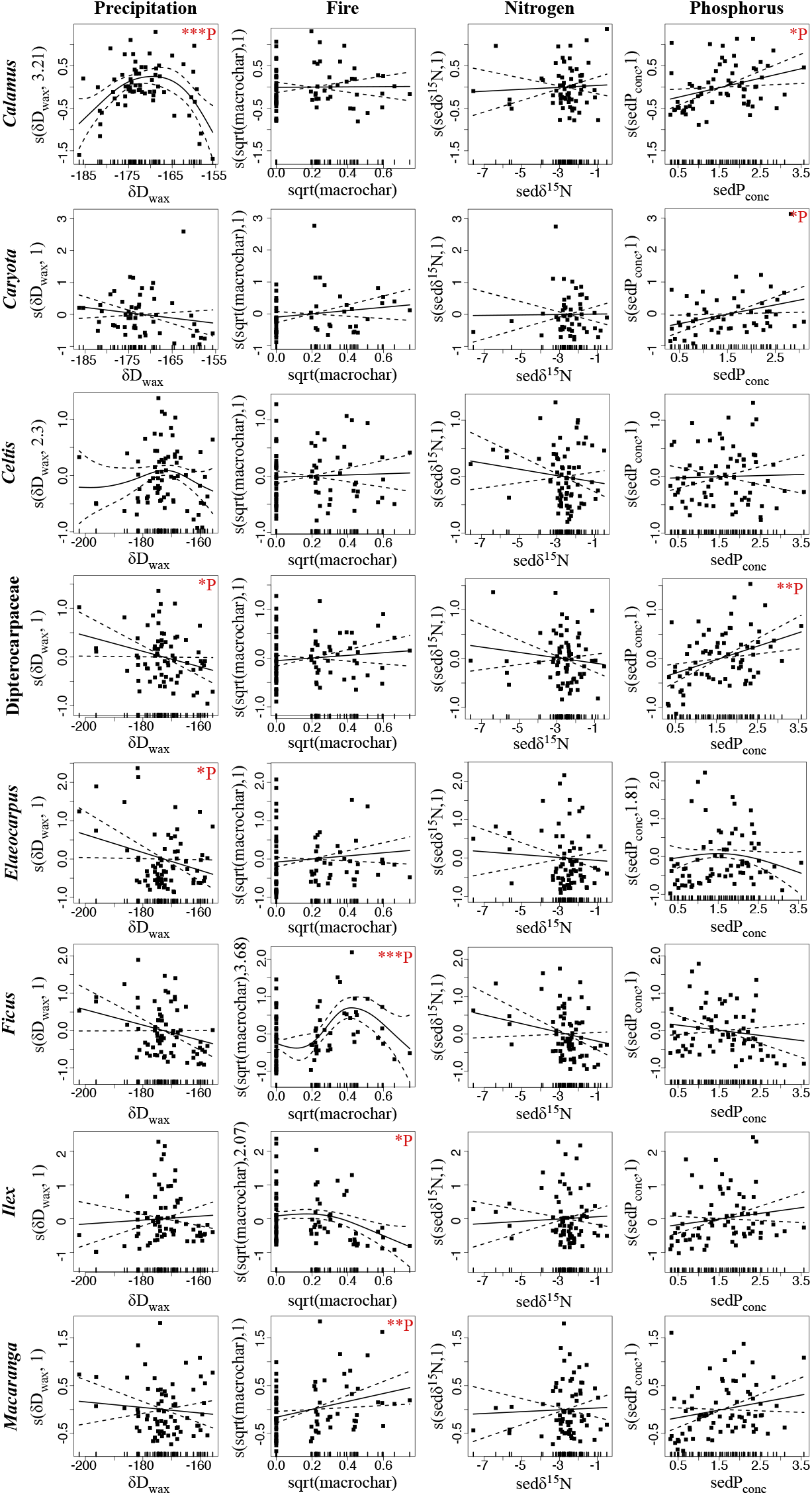
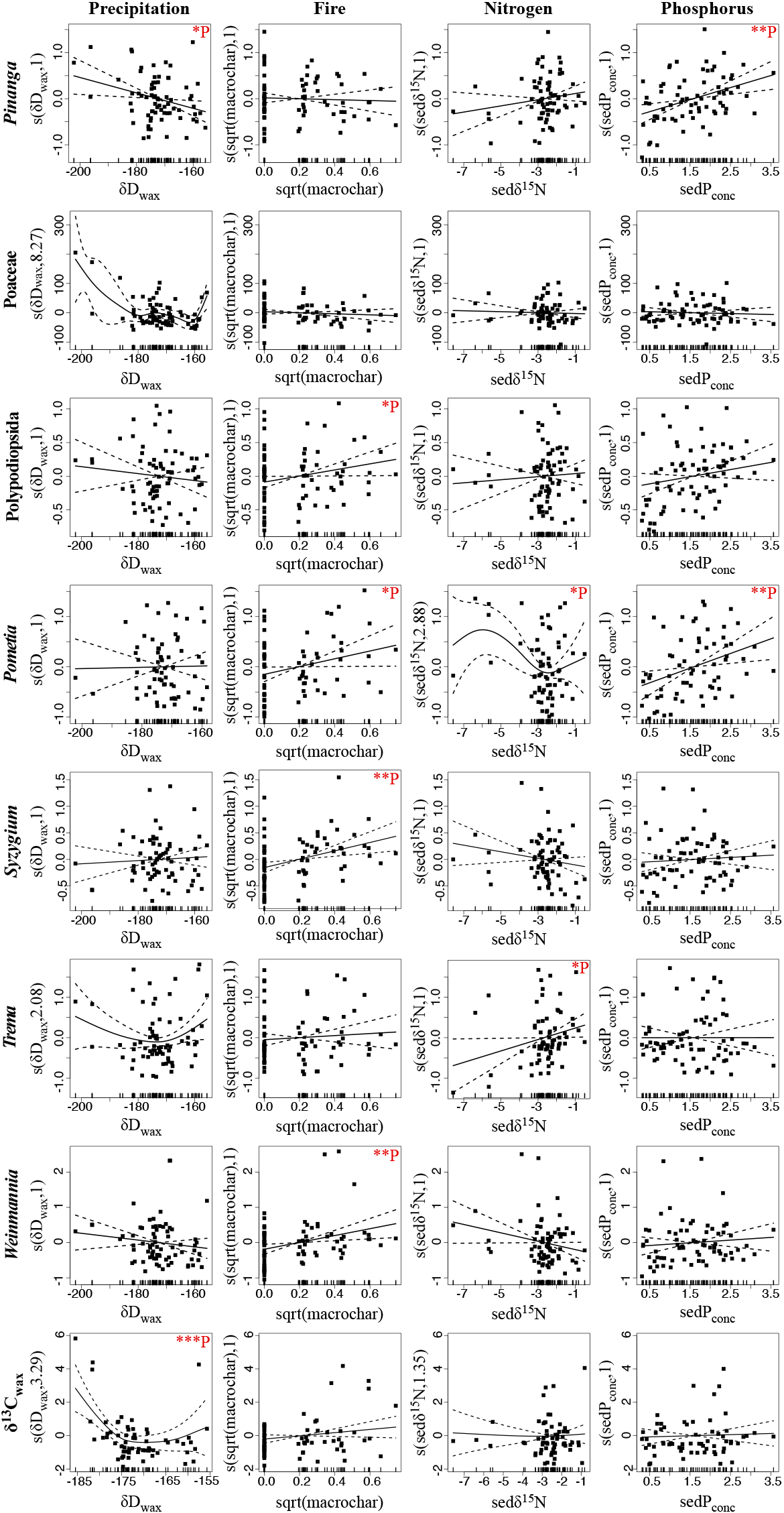
Modelled effects of abiotic factors on plant taxa abundance and C3/ C4 vegetation composition at Bulusan. Plots of GAM smooth functions from the best fitted models for describing the changes in individual taxa’s pollen accumulation rates (proxy for taxa abundance) and δ^13^C values of the *n*C_29_ alkane (proxy for C3/ C4 vegetation composition). General model formula: PAR ~ s(δD_wax_) + s(sqrt(macrochar)) + s(sedδ^15^N) + s(sedP_conc_), distribution family = Gamma, smoothing parameter estimation method = Restricted Maximum Likelihood; and δ^13^C_wax_ ~ s(δD_wax_) + s(sqrt(macrochar)) + s(sedδ^15^N) + s(sedP_conc_), distribution family = Gaussian, smoothing parameter estimation method = Restricted Maximum Likelihood. PARs = pollen accumulation rates (grains cm^−2^yr^−1^), macro = macrocharcoal influx rates (particles m^−2^yr^−1^), δ^13^C_wax_ = δ^13^C values of the *n*C_29_ alkane (‰), δD_wax_ = δD values of the *n*C_29_ alkane (‰), sedδ^15^N = δ^15^N values of bulk sediment (‰), sedP_conc_ = phosphorus concentration of bulk sediment (XRF counts per second). The tick marks on the x-axis are observed data points. The y-axis represents the partial effect of each variable. The dotted lines indicate the 95% confidence intervals. Asterisks denote significance levels: *P-value ≤ 0.05; **P-value ≤ 0.01; ***P-value ≤ 0.001. Data was analysed and plotted using the mgcv 1.8-31 package (Wood 2018) in R version 3.6.3 (R Core Team 2020).

Our results are in accordance with observational studies that found Dipterocarp trees to have particularly low tolerance to drought. For instance, (Nakagawa *et al*. 2010) found that during drought events the mortality of Dipterocarp species can increase up to 30 times, as opposed to 2-11 times for other tree families. Previous research also has established a positive link between ENSO conditions and an Dipterocarp flower production; stronger El Niño drought events usually result in stronger general flowering events where the period between two mass flowering events can be as short as 2 years (Sakai *et al*. 2006). Furthermore, Kaars *et al*. (2009) showed that the higher frequency of El Niño events over the last 250 years was accompanied by higher concentration of Dipterocarpaceae pollen in the marine core from Kau Bay, Indonesia. However, our results indicate that while enhanced ENSO conditions in this region over the past 1,400 years may have led to intensification of mass flowering events, any positive effects of such climatic conditions on pollen production had been surpassed by its negative effects, such as increased tree mortality.

Surprisingly, we found no evidence that Dipterocarpaceae PARs were significantly affected by the changes in the fossil macrocharcoal influx rates (the best-fitted GAM yielded a non-significant effect (p-value >0.05), Table 1), suggesting lack of relationship between Dipterocarp abundance and fire activity. This indicates that the mechanisms which led to the (mass) flowering of mature Dipterocarp trees were uninhibited during the periods when fire activity was higher. Most Dipterocarp species belong to the largest tree category (i.e., >80 cm d.b.h.) and observation studies suggested that such large trees are rarely affected by fire (VAN Nieuwstadt, Van Nieuwstadt, and Sheil 2005). Our results support this understanding.

The positive relationship between sedimentary P_conc_ and Dipterocarp PARs in the Bulusan record suggests a long-term phosphorus limitation to the growth and reproduction in Dipterocarps. Banin *et al*. (2014) found that Dipterocarp trees gain woody biomass faster than otherwise equivalent, neighbouring non-dipterocarp trees, demonstrating that Dipterocarp growth is an important contributor to the overall woody biomass production in Dipterocarp forests (Paoli *et al*. 2008). Though experimental work suggests that the effect of phosphorus availability on the rate of woody biomass increase in Dipterocarps is relatively small on an annual scale (Mirmanto *et al*. 1999), it is possible that this effect compounds on multidecadal scales, especially for large trees. For instance, observational studies of biomass production in Dipterocarp forest plots across nutrient gradients showed that tree biomass increment is positively related to nutrient availability, particularly phosphorus, and that this increment is more strongly related for large trees (Paoli *et al*. 2005; Paoli & Curran 2007; Paoli *et al*. 2008).

On the other hand, the positive relationship between PARs and P_conc_ observed in the Bulusan record could be, at least partly, related to pollen productivity rates. Previous studies have shown that in any mass flowering event, only a portion of reproductively mature Dipterocarp individuals actually produce flowers (Ashton *et al*. 1988; Sakai *et al*. 1999), and this was attributed to the nutritional status of the trees. Ichie *et al*. (2005) showed the high phosphorus cost of mass flowering and subsequent mast fruiting for Dipterocarp species (Ichie *et al*. 2005; Ichie & Nakagawa 2013). Therefore, accumulation of phosphorus has been put forward as a potentially decisive factor in the magnitude and frequency of Dipterocarp reproduction events on the prevailingly nutrient poor soils of Southeast Asia (Ichie & Nakagawa 2013). Given this evidence, decreasing phosphorus availability at the Bulusan site could have potentially decreased the frequency and/ or magnitude of mass flowering in Dipterocarps, resulting in lower production rates of their pollen. Since Dipterocarp pollen production rates may be heavily dependent not only on the tree abundance, but also on climatically mediated flowering rates, future research would benefit from using independent proxies for Dipterocarp tree abundance, such ancient DNA or taxa specific biomarkers, to separate the two signals.

### Influence of climate and nutrient availability on other Dipterocarp forest taxa

Other plant taxa show a wide variety of relationships to the climatic and nutritional conditions at Bulusan (Table 1 and Fig. 3, Table S1). We find a significant negative relationship between the δD_wax_ and the PARs of *Elaeocarpus and significant varying relationship,Pinanga* (Table 1 and Fig. 3, Extended Table S1), suggesting a detrimental impact of drought on the abundance of these taxa. Interestingly, *Calamus* PARs exhibit unimodal relationship to the δD_wax_, indicating that while moderate drought conditions have a positive effect on *Calamus* abundance, the opposite is the case at more extreme drought conditions. Interestingly, none of the taxa exhibited an altogether positive relationship to δD_wax_, suggesting a predominantly negative effect of drier conditions on the Dipterocarp forest vegetation.

Furthermore, the PARs of *Macaranga, Pometia*, Polypodiopsida, *Syzygium* and *Weinmannia* show a significant positive relationship to macrocharcoal influx rates. The apparent positive effect of fire activity on pioneers such as *Macaranga*, *Weinmannia* and Polypodiopsida at our site is in accordance with a number of observational studies which have reported positive effects of fire on the establishment and growth of pioneer plants (Whitmore 1988; Woods 1989; Schulte & Schöne 1996). One explanation of this relationship is that fire removes the pre-existing seedlings of climax taxa from forest gaps, thereby exposing bare soil and facilitating germination and growth of pioneer taxa (Woods 1989; Ashton & Seidler 2014). On the other hand, the surprising lack of negative relationship between PARs and fossil macrocharcoal influx rates among other Bulusan plant taxa (with the exception of *Ilex*) may be partly explained by a limited fire activity at Bulusan Lake and its wider area, as suggested by the relatively low macrocharcoal and microcharcoal counts, respectively (Fig. S2). Consequently, fire events at Bulusan may have not been severe enough to stifle growth and reproduction of plant taxa with a low tolerance to fire.

While PARs of nearly all taxa in the Bulusan record exhibited no significant relationship to nitrogen, the PARs of one third of taxa showed a positive relationship to P_conc_ (*Calamus, Caryota, Pinanga, Pometia*, along with Dipterocarpaceae), providing further support to phosphorus being a major limiting nutrient in Dipterocarp forests.

### Influence of climate and nutrient availability on forest vegetation cover

Our results show a unimodal relationship between δ^13^C_wax_ and δD_wax_ values (Fig. 3b), where the relationship is positive at the lower and higher ends of δD_wax_ values (wet and dry conditions) and negative at intermediate δD_wax_ values (moderate conditions). This indicates that moderate conditions had a positive effect on C3 vegetation (trees and shrubs) but that extremely wet and dry conditions lead to an increase in C4 component (grasses). While the negative effect of extremely wet conditions may seem counterintuitive at first, it could be explained by the dependence of Dipterocarp forest reproduction on ENSO-induced drought events; low frequency of such events may have led to reduced reproduction of mass flowering tree taxa, which in long-term translated to reduced abundance of those taxa, and hence a relatively higher proportion of grasses (though still a low proportion considering the absolute δ^13^C_wax_ values in the Bulusan record never approach the values characteristic of vegetation cover dominated by C4 plants). Nevertheless, the positive relationship between δ^13^C_wax_ and δD_wax_ at the high end of δD_wax_ values highlights the negative effect of more dry conditions on Dipterocarp forest vegetation, possibly through an increased mortality of large trees and subsequent opening of the canopy that favours establishment of grasses.

We found no significant effect of fossil macrocharcoal influx rates on δ^13^C_wax_, despite the overlap of the peaks in both records around 1500 AD (Fig. 2b, t). However, the generally low fossil macrocharcoal influx rates at Bulusan over the last 1,400 years (apart from the 1500 AD event), suggestive of a low fire activity in the area, may explain this apparent lack of relationship.

### Implications for conservation and management

Our findings have implications for the conservation and management of Dipterocarp forests under future climate change. Projected drought risk in 1.5°C and 2°C warmer climate scenarios show robust drying in the mean state across many regions by the end of the 21st century, including Southeast Asia (Lehner *et al*. 2017; Cook *et al*. 2020). Our results suggest that a stronger drought regime could have profound impacts on the composition, structure and functioning of Dipterocarp forests, primarily through a negative impact on the abundance of Dipterocarp trees. As giant, old-growth trees, with height over 70 meters, a very complex canopy, and wide base forming an extensive buttressing, Dipterocarp trees embody the concept of tropical mega-trees (Pinho *et al*. 2020). Recent studies highlighted the unique ecological roles of mega-trees such as building the emergent forest layer, providing unique microhabitats inside the complex canopy, and acting as a keystone source of food and nesting sites. Furthermore, tropical mega-trees are disproportionately large contributors to forest productivity, above ground biomass and carbon stocks (Slik *et al*. 2013; Stephenson *et al*. 2014; Lutz *et al*. 2018). The importance of Dipterocarp trees for the ecological integrity of Dipterocarp forests is further amplified through their dominance (Appanah 1985, Paoli *et al*. Slik 2008) and mass flowering behaviour (Burgess 1972; Cockburn 1975; Appanah 1985, Sakai *et al*. 1999), which profoundly shape pollination and food production patterns in these forests. While it has been hypothesized that a stronger drought regime may also increase the frequency and strength of mass flowering, our results suggest that a positive effect of drought is unlikely to offset its negative impacts, e.g. increased tree mortality. Given the vulnerability of Dipterocarps to heightened drought and the projected increase in the frequency and intensity of drought in their geographic range (Amnuaylojaroen & Chanvichit 2019; Pörtner *et al*. 2022), along with endangered status of most Dipterocarp species (Bartholomew *et al*. 2021), considerable efforts need to invested into ensuring sustainable land use practices in the remaining Dipterocarp forest tracts.

The last few decades have also seen a marked increase in the frequency and magnitude of fire events in Dipterocap forests (Salafsky 1994; Folkins *et al*. 1997; Salafsky 1998; Siegert *et al*. 2001; Huijnen *et al*. 2016). The observed trend has been linked to the compounding effects of more severe El Niño events (van der Werf *et al*. 2004; Wooster *et al*. 2012; Chen *et al*. 2017; Nurdiati *et al*. 2022) and anthropogenic activities such as forest fragmentation and degradation (Curran *et al*. 1999; Chapman *et al*. 2020). Therefore, it is highly likely that a future intensification of ENSO coupled with anthropogenic activities will result in heightened fire activity in the region. While our study suggests that none of the considered plant taxa have been negatively affected by fire activity over the last 1,400 years, our fossil macrocharcoal record suggests an overall limited fire activity at Bulusan during this time period. Notably, the single period of markedly higher fire activity in the Bulusan record around 1500 AD closely overlaps with a significant increase in the C4 vegetation. Hence, our results do not exclude the possibility that prolonged periods of higher fire activity would not lead to larger prevalence of plant pioneers in forest floor gaps created by burning, increased canopy openness and a reduced above ground biomass, as suggested previously by observational studies (Whitmore 1988; Ghazoul 2016). Given the severity of the synergistic effects that drought and fire can have on Dipterocarp forests, strategies for minimising fire occurrence and magnitude, particularly during droughts, should be prioritised to prevent further degradation of these globally important forests.

Our understanding of the nutrient cycles that underlie the productivity of tropical rainforests worldwide remains limited. The prevalent view of primary tropical rainforests is that they inhabit infertile soils, where most of the nutrients are tied in the living matter, and their high production relies on fast and efficient nutrient cycling (Cleveland *et al*. 2011). However, our study shows that the availability of key limiting nutrients, namely phosphorus, to tropical forest plants can vary considerably on multidecadal to centennial scales (in line with studies on nitrogen availability in higher latitudes e.g. (Jeffers *et al*. 2011, 2012; McLauchlan *et al*. 2017) and that this variation affects various plant taxa including Dipterocarp trees, thus possibly playing a key role in shaping the productivity of Dipterocarp forests. This finding raises important questions for the conservation and management of Dipterocarp forests that can be further tested with palaeoecological data. For instance, if phosphorus availability has a considerably positive influence on the growth or reproduction of large tropical trees, then all else being equal, are tropical rainforests growing on more fertile soils more resilient to drought and other environmental perturbations on longer time scales? Our study is the first to link changes in pollen accumulation rates to the changes in availability of phosphorus spanning hundreds to thousands of years, demonstrating the potential of paleo-records for studying the effects of nutrient limitation on the tropical rainforest biome. This knowledge could vastly improve current models of global climate-vegetation dynamics under warming scenarios given the central role of this biome in regulating Earth’s climate and biogeochemical cycles (Brown & Lugo 1982; Melillo *et al*. 1993; Cleveland *et al*. 1999; Bonan 2008; Fleischer *et al*. 2019).

While our findings highlight the great value of palaeoecological data for advancing our understanding of the impacts of climate and nutrient availability on Dipterocarp forests, additional records are needed to test the generality of the findings reported here across larger temporal and spatial scales in order to improve the forecasts of tropical rainforest dynamics in a warming world.

## Data availability

All data needed to appraise the conclusions in the paper are given in the Main Text and the Supporting Information. Any additional data related to this paper may be requested from the authors.

## General

We thank the Philippine Bureau of Fisheries and Aquatic Resources and the Philippine Department of Environment and Natural Resources, as well as the Bulusan Volcano Natural Park staff, for permitting and facilitating our research at the Bulusan Lake. We also thank Professor Keith Bennett (University of St Andrews) and Philip Bartilet (Bulusan Volcano Natural Park) for fieldwork support, and Iris Van Der Veen (University of Potsdam) and Bernd Meese (Fraunhofer Institute for Manufacturing Engineering and Automation) and Anne Thoisen (University of Aarhus) for laboratory assistance. For assistance with the figures we thank Marwan Butrous.

## Author contribution

AP initiated and designed the study and conducted the fieldwork. AP planned the sampling scheme and led the generation and analysis of data with assistance from AWRS, AS, OR and DS. AP, KJW and DS acquired financial support. All authors participated in interpreting the results and writing the manuscript in the lead of AP.

## The following Supporting Information is available for this article

**Fig. S1** Age-depth model for the Bulusan sedimentary sequence.

**Fig. S2** Microcharcoal influx rates in relation to macrocharcoal influx rates in the Bulusan record.

**Fig. S3** Pollen diagram of the sedimentary sequence from Bulusan Lake.

**Fig. S4** Comparison of the records of elements concentrations, magnetic susceptibility and bulk sediment phosphate oxygen isotopes from the Bulusan sedimentary sequence.

**Fig. S5** Principal Component Analysis results of the abundance dynamics of main geochemical elements and the magnitude of magnetic susceptibility in the Bulusan sedimentary sequence.

**Fig. S6** Comparison of the records of bulk sediment phosphorus concentrations, δDwax values, macrocharcoal influx rates, and Ba/Ti ratio from the Bulusan sedimentary sequence.

**Fig. S7** Significant periods of change in the key paleo-records from Bulusan Lake derived from the best fitted generalised additive model trends.

**Fig. S8** Modelled effects of abiotic factors on plant taxa abundance at Bulusan using generalised additive models with a Quasi-Poisson distribution. Plots of GAM smooth functions from the best fitted models.

**Table S1** AMS14C dates of the Bulusan sedimentary sequence.

**Table S2** Summary of results of the best fitted generalised additive models with a Quasi-Poisson distribution for describing the effects of abiotic factors on plant taxa abundance at Bulusan.

**Table S3** Description of the best fitted generalised additive models of temporal dynamics of paleo-records from Bulusan Lake with GCV-based smoothness selection.

**Table S4** Description of the best fitted generalised additive models of temporal dynamics of paleo-records from Bulusan Lake with a continuous-time AR(1) process estimated using REML smoothness selection.

**Table S5** Description of the best fitted generalised additive models with a Gamma distribution for describing the effects of abiotic factors on plant taxa abundance and C3/ C4 vegetation composition at Bulusan.

**Table S6** Description of the best fitted generalised additive models with a Quasi-Poisson distribution for describing the effects of abiotic factors on plant taxa abundance at Bulusan.

**Methods S1** Age-depth modelling.

**Methods S2** Magnetic susceptibility records.

**Methods S3** Microcharcoal influx rates.

**Methods S4** Phosphate oxygen isotope analysis.

**Notes S1** Potential drivers of charcoal influx rates in the Bulusan record.

**Notes S2** Itrax XRF and magnetic susceptibility records.

**Notes S3** Potential drivers of changes in phosphorus availability at Bulusan.

**Notes S4** Pollen accumulation rates as a proxy for Dipterocarp dynamics.

**Notes S5** Identifying significant periods of change with GAMs.

**Notes S6** Comparison of GAMs with Gamma and Quasi-Poisson distribution.

