## Supporting Information for "Long-Term Ecological Responses of a Dipterocarp Forest to Climate Changes and Nutrient Availability"

Article acceptance date:

The following Supporting Information is available for this article

**Fig. S1** Age-depth model for the Bulusan sedimentary sequence.

**Fig. S2** Microcharcoal influx rates in relation to macrocharcoal influx rates in the Bulusan record.

**Fig. S3** Pollen diagram of the sedimentary sequence from Bulusan Lake.

**Fig. S4** Comparison of the records of elements concentrations, magnetic susceptibility and bulk sediment phosphate oxygen isotopes from the Bulusan sedimentary sequence.

**Fig. S5** Principal Component Analysis results of the abundance dynamics of main geochemical elements and the magnitude of magnetic susceptibility in the Bulusan sedimentary sequence.

**Fig. S6** Comparison of the records of bulk sediment phosphorus concentrations, $\delta D_{wax}$ values, macrocharcoal influx rates, and Ba/Ti ratio from the Bulusan sedimentary sequence.

**Fig. S7** Significant periods of change in the key paleo-records from Bulusan Lake derived from the best fitted generalised additive model trends.

**Fig. S8** Modelled effects of abiotic factors on plant taxa abundance at Bulusan using generalised additive models with a Quasi-Poisson distribution. Plots of GAM smooth functions from the best fitted models.

**Table S1** AMS14C dates of the Bulusan sedimentary sequence.

**Table S2** Summary of results of the best fitted generalised additive models with a Quasi-Poisson distribution for describing the effects of abiotic factors on plant taxa abundance at Bulusan.

**Table S3** Description of the best fitted generalised additive models of temporal dynamics of paleo-records from Bulusan Lake with GCV-based smoothness selection.

**Table S4** Description of the best fitted generalised additive models of temporal dynamics of paleo-records from Bulusan Lake with a continuous-time AR(1) process estimated using

REML smoothness selection.

**Table S5** Description of the best fitted generalised additive models with a Gamma distribution for describing the effects of abiotic factors on plant taxa abundance and C3/ C4 vegetation composition at Bulusan.

**Table S6** Description of the best fitted generalised additive models with a Quasi-Poisson distribution for describing the effects of abiotic factors on plant taxa abundance at Bulusan.

**Methods S1** Age-depth modelling

**Methods S2** Magnetic susceptibility records

**Methods S3** Microcharcoal influx rates

**Methods S4** Phosphate oxygen isotope analysis

**Notes S1** Potential drivers of charcoal influx rates in the Bulusan record

**Notes S2** Itrax XRF and magnetic susceptibility records

**Notes S3** Potential drivers of changes in phosphorus availability at Bulusan

**Notes S4** Pollen accumulation rates as a proxy for Dipterocarp dynamics

**Notes S5** Identifying significant periods of change with GAMs

**Notes S6** Comparison of GAMs with Gamma and Quasi-Poisson distribution

**Methods S1** Age-depth modelling

A high-resolution age-depth model was established for the Bulusan sedimentary sequence using 10 AMS$^{14}$C dates measured at the NERC Radiocarbon Facility, East Kilbride (Table S1) and the surface sediment age set to 2013. The radiocarbon analysis was undertaken on terrestrial plant macrofossils (e.g., leaves, twigs) to avoid reservoir effects and minimise age uncertainty. The model was computed with the Bacon 2.2 package at 1 cm resolution (Blaauw & Andrés Christen 2011, 2015) using R version 3.2.4 (R Core Team 2020) and the IntCal13 radiocarbon calibration curve for the Northern Hemisphere (Reimer *et al.* 2013) (Fig. S1, from Prohaska *et al.* (2022). According to the model, the resolution of individual sediment samples is ~20 years.

**Methods S2** Magnetic susceptibility records

Magnetic susceptibility was measured on intact Bulusan cores using the Barrington MS2 Magnetic Susceptibility System made of a magnetic susceptibility meter coupled with a core logging sensor at 2 cm spatial resolution and 11s SI (9s CGS) measurement period.

**Methods S3** Microcharcoal influx rates

Microcharcoal influx rates (particles $cm^{-2}yr^{-1}$) were calculated by counting the total number of particles in each sample, determining its concentration using known marker concentrations (Stockmarr 1971) and dividing these with the corresponding sediment accumulation rate (Clark 1988; Mooney & Tinner 2010).

**Methods S4** Phosphate oxygen isotope analysis

Sediment samples were sequentially extracted following the protocol of Amelung *et al.* (2015); 2-8 g of dried sediment (depending on phosphorus (P) concentration and sample availability) was extracted with 0.5 mol NaHCO3 at a soil to solution ratio of 1:15. After 16-

hour shaking, the samples were centrifuged, supernatant disposed of and the solid sample re-extracted with 0.1 mol NaOH at soil to solution ratio of 1:15. After another 16-hour shaking the procedure to remove the supernatant was repeated, sediments were finally extracted in 40 ml of 1 mol HCl. The HCl extract was purified to silver phosphate following the well established protocols of Tamburini *et al.* (2010). Oxygen isotope analysis of silver phosphate ($\delta^{18}O$-P) was undertaken on an Elementar Pyrocube elemental analyser (EA) coupled to a Geovision Isotope Ratio Mass Spectrometer (IRMS). 200-300 µg of silver phosphate was weighed into high purity silver caps and introduced to the EA via the built-in autosampler in a flow of lab grade (99.9%) He. The sample was pyrolysed at 1450 ºC in the presence of graphite, carbon black and glassy carbon, where oxygen from phosphate combines with available carbon forming CO gas which is introduced into the IRMS Raw isotope values are corrected using the Ionos software for blank and drift issues, if necessary, before correction using a two point calibration, based on the externally calibrated values of international standards USGS 80 and B2207, with Alfa 2 (in house standard), acting as a check standard. Oxygen isotope values are corrected to the international VSMOW scale. Typical precision is <0.3‰ for $\delta^{18}O$-P based on within run replication of reference materials.

**Notes S1** Potential drivers of charcoal influx rates in the Bulusan record

Macrocharcoal influx rates in the Bulusan record ranged between 0-1.8 particles $cm^{-2}yr^{-1}$ (Fig. 2b). These are low values even when compared to other records from tropical rainforests, which document maximum macrocharcoal influx rates from 500 to up to 4000 particles $cm^{-2}yr^{-1}$ (e.g., Haberle (2005); Hermanowski *et al.* (2015)) and maximum macrocharcoal concentrations from 100 to over 4000 particles $cm^{-3}$ (e.g., Biagioni *et al.* (2015); Cole *et al.* (2015)). Similarly, microcharcoal influx rates were consistently low along the Bulusan sequence (i.e., 0-22 particles $cm^{-2}yr^{-}1$, Fig. S2) in comparison to other lacustrine cores from tropical forest regions which record a maximum of 40,000-80,000 particles $cm^{-2}yr^{-1}$ (e.g., Haberle (2005); Cordeiro *et al.* (2014)). The low charcoal counts at Bulusan site have precluded calculations of fire frequency as the signal-to-noise index (SNI) values are lower than 1 for most of the record, indicating it is not appropriate for peak analysis. Instead, the overall charcoal accumulation can be interpreted as a general proxy for fire activity: rate biomass burning, which integrates both rate of burning, and biomass burned per fire (Marlon *et al.* 2008; Marlon *et al.* 2013; Marlon *et al.* 2016).

Given the relatively low charcoal influx rates in the Bulusan record, it is difficult to pinpoint key drivers of fire activity at the site. Overall, the charcoal influx rates are low enough to suggest fires of climatic origin, such as ENSO-induced drought conditions. Previous work has demonstrated the positive relationship between ENSO-induced drought and fire (Fuller & Murphy 2006; Langner & Siegert 2009; Wooster *et al.* 2012), and while undisturbed tropical rainforests are usually highly resistant to fire due to low amounts of burning fuel, the association of fire with severe ENSO-induced droughts have been recorded as early as 1914 in the Southeast Asian region (Goldammer *et al.* 1996). However, the limited overlap between ENSO-induced drought conditions and enhanced fire activity in the Bulusan record suggests that factors other than ENSO dynamics have also influenced fire dynamics at the site. Previous studies from other regions have demonstrated that volcanic activity can cause widespread fires that last several years to several decades (Wilmshurst & McGlone 1996; Zhao *et al.* 2015). Still, the period of enhanced fire activity in the Bulusan record does not coincide well with tephra layers, suggesting a limited effect of volcanic activity on fire dynamics at the Bulusan lake. Humid tropical environments have low incidence of natural fire (Scott 2000) and therefore high charcoal influx rates in absence of drier conditions in tropical rainforest records are often linked to humans (e.g., Bush *et al.* (2008)). However, the relatively low charcoal rates at the Bulusan site fail to provide compelling evidence of human activities. Furthermore, while early written accounts of human settlement in the wider Bulusan area can be traced back to AD 1630, when it was declared a parish independent from Casiguran, and excavated artefacts from native burial grounds document a number of human settlements in this area prior to Spanish arrival (Abada Gatumbato *et al.* 2011), no archaeological evidence has been found in the close vicinity of the lake to date. Though low microcharcoal counts during the peak in macrocharcoal influx rates at ~ 1500 AD suggest that this was a period of localised fire at the site, such as those caused by human-activities (Bennett *et al.* 1990), more charcoal records from neighbouring areas are needed to confirm the lack of spatial and temporal autocorrelation of fire activity on the wider landscape characteristic of human burning. Notably, the decrease in macrocharcoal influx rates from its peak values at ~1500 AD to present conflicts with the narrative of human-related fire activity at the site, as it is unlikely that human burning would have reduced towards present times given the accelerating expansion of agriculture and urbanisation in the wider area. Finally, an increase in Poaceae pollen in humid tropics can also be indicative of human disturbance (Bush 2002), yet Poaceae PARs remained relatively low throughout the Bulusan record apart from more recent times, and with no significant increases even during periods of increased fire activity. Given these different lines of evidence, humans are unlikely to be a prominent driver of fire activity at the site, though they may have contributed to some extent to the observed fire patterns.

**Notes S2** Itrax XRF and magnetic susceptibility records

The XRF results show relatively high counts and low noise-to-signal ratio for some elements in the Bulusan records, such as K and Ca (Fig. S4). In contrast, other elements such as Si, Sr and P have relatively low counts and high noise-to-signal ratio. Nevertheless, pronounced peaks and troughs can be observed in nearly all of the element records.

Magnetic susceptibility values are relatively low throughout the Bulusan sequence, except for the peaks that overlap with the tephra layers. These results demonstrate a low proportion of ferromagnetic material in the Bulusan sediments between volcanic eruptions. The generated magnetic susceptibility record has a relatively low noise-to-signal ratio.

**Notes S3** Potential drivers of changes in phosphorus availability at Bulusan

Results from the XRF scanning analysis show distinct fluctuations in bulk sediment phosphorus concentrations ($sedP_{conc}$ hereafter) in the Bulusan sedimentary sequence over the last 1,400 years. There are several potential drivers of P variability in lake sediments including rock weathering (Ruttenberg 2003), soil leaching and erosion (Ruttenberg 2003), sediment reworking (Robbins 1982), animal-mediated translocation of nutrients (Doughty *et al.* 2013) and changes in soil pH in the lake's watershed (Broberg & Persson 1984; Jansson *et al.* 1986; Kopáček *et al.* 2015).

Weathering of continental bedrock is the main source of P to the soils that support continental vegetation over geological time scales (i.e., $10^3$s to $10^6$s years). P is weathered from bedrock by dissolution of P-bearing minerals e.g., apatite by naturally occurring acids, namely from microbial activity (e.g.  (Cosgrove 1977; Frossard *et al.* 1995). Changes in the rates of tectonic uplift and exposure of P-bearing rocks can therefore cause changes in the P availability in an ecosystem which would be eventually reflected in its lake sediments.

Soil development can also affect the deposition of P in lake sediments on longer time scales (i.e., $10^3$s to $10^4$s years). According to the classic model by Walker and Syers (1976), later validated by numerous studies (e.g. Smeck (1985); Crews *et al.* (1995); Filippelli & Souch (1999)), soil development entails systematic changes in the total amount and chemical form of P; while initial stages have P present mainly as primary minerals (e.g. apatite), its fraction decreases in mid-stage soils which instead harbour increasing levels of less-soluble secondary minerals and organic P, until finally in late stages of soil development P is partitioned primarily between refractory minerals and organic P. Leaching is the process by which water-soluble plant nutrients such as P are lost from the soil, by means of rivers, rain or irrigation (agriculture). Changes in river discharge, precipitation amount, fire activity or agricultural practices can influence the leaching rates of soluble P from the soil and its deposition in lake sediments on shorter time scales (i.e., annual to millennial scales) (Schlesinger 1997; Ruttenberg 2003). Similarly, changes in the soil erosion rates, due to precipitation variability, landslides or changes in land-use can affect the influx of insoluble P, bound by various soil constituents, to the lake.

Sediment reworking is a mechanism by which P abundance can vary in lake sediments independently of its availability on the land. For instance, rates of sediment reworking by high concentration of zoobenthos (e.g., oligochaete worms) in North American Great Lakes are comparable to or exceed sedimentation rates (Robbins 1982). Thus, significant changes in the concentration of zoobenthos species over time can alter P sedimentation rates following its introduction to the lake. Furthermore, changes in the redox-driven coupled Fe–P cycling of bottom waters and benthic P efflux can also affect P abundance in the sediments (Mortimer 1941; Mortimer 1942; Boström *et al.* 1988). Microbial reduction and reoxidation/ re-precipitation of ferric oxyhydroxides controls P release and retention rates by sediments through both direct and indirect responses of microbial community to changes in redox state of bottom waters (e.g., Mortimer (1941), Mortimer (1942), Gächter & Meyer (1993)).

Animals are an important vector of nutrients in terrestrial ecosystems, and large bodied animals play a central role in nutrient translocation because they travel larger distances and digest food at a slower rate in comparison to smaller animals (Demment & Van Soest 1985; Kelt & Van Vuren 2001). For instance, previous work has suggested that reduction in the biomass of large mammals due to Pleistocene megafaunal extinctions has significantly reduced the lateral flux of P on all continents outside of Africa (Doughty *et al.* 2013). Given that permanent water bodies are known hotspots of animal activity in forested landscapes, changes in the biomass of large mammals in such landscapes may lead to notable change in P deposition in lacustrine sediments.

Finally, changes in soil pH in a lake's watershed can also affect the influx rates of nutrients to the lake (Broberg & Persson 1984; Jansson *et al.* 1986; Kopáček *et al.* 2015). P is particularly affected by the soil pH and its solubility is usually highest in neutral soils (i.e., pH 6.5 -7.5). At alkaline pH values (pH >7.5) phosphate ions tend to react quickly with calcium (Ca) and magnesium (Mg) to form less soluble compounds, while at acidic pH values (pH <6.5), phosphate ions react with aluminium (Al) and iron (Fe) to again form less soluble compounds (Lajtha & Harrison 1995). Soil acidification by means of volcanic ash deposition has been well documented (Cronin *et al.* 1998; Cronin *et al.* 2003; Ranatunga *et al.* 2009; Wilson *et al.* 2011), suggesting that changes in volcanic activity over time can exert significant changes in P solubility, and consequently, its availability to plants as well as the rate of its leaching out of the soil and into the lake.

To determine the main driver(s) behind the variability of P abundance in the Bulusan sediments, we undertook the following analyses. Firstly, we performed a Principal Component Analysis (PCA) on the chemical elements detected by the Itrax XRF Core Scanner to test the relationship between P and the main leachable elements of volcanic ash material, i.e., silicon (Si), potassium (K), calcium (Ca) and strontium (Sr). Prior to the PCA application, Itrax data were Hellinger transformed to reduce the influence of abundant elements. To investigate the effect of soil erosion on the P abundance, we reconstructed the record of organic to inorganic matter ratio using the ratio of barium to titanium (Ba/Ti) values (Croudace *et al.* 2006) and compared it to the $sedP_{conc}$ record. Finally, to examine the influence of weathering and burning on P abundance, we compared the $nC_{29}$ alkane δD values (hereafter $\delta D_{wax}$) and macrocharcoal influx rates record with the $sedP_{conc}$ values.

Analysis of the Bulusan sediments suggests a negative relationship between P and the volcanic ash deposition proxies (Si, K, Ca, Sr) and the magnetic susceptibility (MS) values. $K^+$ and $Ca^+$ cations are among the most abundant ionic species in volcanic ash leaches (Jones & Gislason 2008; Wilson *et al.* 2011), and these leaches also represent an important source of Sr in nature

(Koshikawa *et al.* 2016), making the three chemical elements suitable proxies for volcanic ash deposition. Similarly, magnetic susceptibility has been shown to be particularly useful for detecting the presence of tephra layers in lacustrine sequences because tephra is usually rich in ferromagnetic material, which shows up as peaks in magnetic susceptibility values (Oldfield *et al.* 1983; Oldfield 1988; Lozano-García *et al.* 1993). The PCA analysis on the individual abundances of chemical elements and magnetic susceptibility magnitude in the Bulusan record shows clear clustering of Si, K, Ca, Sr, and MS, demonstrating similar dynamics among these four variables and suggesting a common source (Fig. S5). We argue that these four variables capture the signal of volcanic ash deposition at the Bulusan site over the last 1,400 years. In contrast, K, Ca, Sr, and MS are separated from other elements, including P, Al, and Fe, by the first PCA axis, suggesting that the latter have not been introduced into the sediments by ash deposition. Fig. S4 shows that the peaks in Si, K, Ca, Sr, and MS match the troughs in the P abundance and vice versa during the 550-1050 AD period, and a gradual but continuous growth in P values takes place during the prolonged period of low ash deposition between 1050-1850 AD.

These findings point towards P solubility via volcanic ash acidification of the soil as the most likely driver of P fluctuations recorded in the Bulusan sediments. The proximity of the active Mt Bulusan volcano (i.e., ~4 km up slope from the lake), whose most frequent historic activities have been acidic silica-rich ash-fall eruptions (Delfin *et al.* 1993; McDermott *et al.* 2005; Siringan *et al.* 2018), further supports this interpretation. The deposition of such volcanic ashes can significantly reduce soil pH, and subsequently, reduce the solubility of P forms (Cronin *et al.* 1998; Cronin *et al.* 2003; Ranatunga *et al.* 2009; Wilson *et al.* 2011), including phosphates (i.e., H2PO4- and HPO42) which are the main source of P to plants (Sutton & Larsen 1964). We therefore hypothesise that during the intervals of volcanic ash deposition in the Bulusan area, the pH of the soils surrounding the Bulusan Lake decreased, and the overall solubility of phosphates reduced, leading to a decrease in their availability to plants and their rain-based transport to the lake.

Additional evidence to support this scenario may be derived from the lake sediment phosphate oxygen isotope record ($\delta^{18}O$-P). Due to P limitation within tropical forests, the demand for and turnover of available P is high. The enzyme driven cycling of P leads to an isotopic resetting of the original $\delta^{18}O$-P, derived from the source P (bedrock), toward an isotope value based on an isotopic equilibrium with temperature and soil water (Chang & Blake 2015). In locations such as Bulusan which have P sources of volcanic origin (bedrock or ash deposits), which have low $\delta^{18}O$-P values globally (Smith *et al.* 2021), this leads to large differences between the original source $\delta^{18}O$-P value and the equilibrium value (Helfenstein *et al.* 2018). The Bulusan $\delta^{18}O$-P record has predominantly high $\delta^{18}O$-P values +20 to +22‰ indicative of isotopic equilibrium, with two distinct deviations to lower isotope values at (700 AD and 925 AD) coinciding with the beginning of peak in trace elements (Si, K, Ca, Sr) concentrations and in magnetic susceptibility signal (Figure S5). These negative isotope deviations indicate a short-lived change in P source, most logically indicative of P originating from ashfall which has a lower volcanic endmember isotope signature. We therefore hypothesise that under an ashfall scenario, in addition to P with an isotope value at or near isotopic equilibrium (+20 to +22‰), additional volcanic P will be introduced (isotopic value of 0 to +10‰). Mineral P derived from ashfall will initially preserve a volcanic endmember isotope signature due to acidic conditions in the soil system. This mineral P will become available to plants slowly over the following years and decades through enzymatic and microbial activity. As it is utilised, this P will then equilibrate and retain an isotope signature related to temperature and soil water $\delta^{18}O$. This can explain why we see in Bulusan sediments a sudden drop in $\delta^{18}O$-P (coinciding with peak values in volcanic trace elements concentrations and magnetic susceptibility) that is followed by a recovery to higher P isotope and $sedP_{conc}$ values over the subsequent decades. Nevertheless, further studies are needed to test this hypothesis, and determine the exact mechanisms underlying the dynamics of P deposition and recycling in the Bulusan area.

Further analysis has yielded little evidence in support of the other potential drivers of P variability in the Bulusan record. Firstly, weathering of continental bedrocks and soil development influence P abundance in soils on geological time scales and they are thus an unlikely cause of the decadal to centennial variability in P abundance recorded in the Bulusan sediments. Secondly, comparison between the $sedP_{conc}$ record and $\delta D_{wax}$ values reveal a stark difference between the two variables (Fig. S6). Of particular importance is a lack of response in P abundance to the large decrease in precipitation between ~1650-1900 AD. Moreover, other palaeo-records from the region suggest relatively stable air temperature conditions over this period, with somewhat colder conditions ~1600-1850 AD (Mann & Jones 2003; Jones & Mann 2004). Furthermore, a poor overlap between the changes in macrocharcoal influx rates and those of $sedP_{conc}$ (Fig. S6), as well as generally low charcoal counts indicative of low fire activity, make it improbable that fire exerted notable influence on the leaching of P at the site. It is hence overall unlikely that differential soil leaching has caused the changes in P abundance recorded in the Bulusan sediments over the last 1,400 years. Similarly, while the Ba/Ti record indicates a large increase in organic matter content in the Bulusan sequence between ~950-1250 AD, this change has a poor overlap with the changes in $sedP_{conc}$ values (Fig. S6), suggesting that the erosion at Lake Bulusan over the last 1,400 years is not a major contributor to the recorded changes in P abundance. Furthermore, the basin from which the sediment cores were collected is more than 12 m deep which minimises the chances of sediment reworking by external forces such as wind and terrestrial animals. Moreover, fine laminations of diatomaceous origin detected throughout the sedimentary sequence provide evidence against sediment reworking by benthic organisms (Fig. 1d), while the different dynamics of P and Fe demonstrated by the PCA analysis (Fig. S5) offer little support towards changes in redox state of bottom waters being a major driver of P abundance in the Bulusan sediments. While algal blooms and die-off can create peaks in nutrient deposition in lake sediments, these processes are dependent on nutrient input from external sources, and planktonic dynamics in the lake would be primarily the consequence rather than the cause of changes in nutrient abundance in lake sediments (Smith 1983; Anderson, Glibert, and Burkholder 2002; Conley *et al.* 2009). Consequently, planktonic activity is an unlikely driver of the recorded $sedP_{conc}$ changes in the Bulusan sequence. Finally, no *Sporormiella* spores were found in the Bulusan record during pollen identification and counting, suggesting the biomass of large mammals at the Bulusan site has been consistently low over this time period, making animal-based translocation of P an improbable driver of the recorded fluctuations in P abundance. Considering the different lines of evidence discussed above, we conclude that volcanic ash deposition was likely the dominant driver of P abundance changes in the Bulusan record over the last 1,400 years.

**Notes S4** Pollen accumulation rates as a proxy for Dipterocarp dynamics

Morphological analysis of fossil pollen is the most commonly used method for reconstructing past plant dynamics. Pollen accumulation rates (PARs) of a particular plant taxon provide independent estimates of its quantity on a landscape. Consequently, PARs have been widely used to reconstruct the dynamics of individual plant taxa over time (Bennett & Willis 2001), including their population size (e.g., Magyari *et al.* (2011); Jeffers *et al.* (2012)), abundance (e.g., Davis (1969); Bjune & Birks (2008)), and biomass (e.g., Mazier *et al.* (2012); Jeffers *et al.* (2015)). However, it is important to consider life history traits of the taxon in question when interpreting its PARs. Dipterocarps are long lived and slow growing species whose size at first flowering is considerably smaller than their maximum size, meaning that there may be a large disparity in the sizes of flowering individuals across the landscape. Hence, Dipterocarp PARs will be influenced both by the number of flowering individuals on a landscape (i.e., absolute abundance) and their size distribution.

Furthermore, previous work has shown that PARs of some plant taxa can be affected by changes in pollen productivity over time, for instance due to changes in environmental conditions (e.g., temperature, precipitation, $CO_2$ concentration) (Mazier *et al.* 2012). Dipterocarp trees in particular have irregular flowering patterns which have been linked to

ENSO-induced drought episodes which take place every 2-7 years. Consequently, changes in flower production rates in response to shifts in environmental conditions could influence Dipterocarp PARs on longer time scales. It is therefore probable that Dipterocarp PARs in the Bulusan sequence reflect both the number and the size of Dipterocarps trees on a landscape and their pollen productivity.

Changes in pollen taphonomy, i.e., depositional conditions of pollen grains, can also influence lacustrine PARs records. For instance, varying water levels can change lake size and shape, and consequently its pollen deposition rate (Bennett and Buck 2016). Similarly, changes in the sedimentation rate due to changing biotic activity or soil erosion intensity can influence the concentration of pollen in the sediments (Giesecke and Fontana 2008). Moreover, alternating sediment types and drying of the sediments can cause changes in the preservation conditions across the sedimentary sequence, especially with cores spanning long time scales, and this in turn can alter pollen concentrations. We have minimised these taphonomic effects through a careful choice of the coring site. Bulusan lake is a permanent, medium-sized lake of considerable depth and steep slopes, and hence any past changes in water levels are unlikely to have brought about major modifications in its size and shape. Because Bulusan sediments are uniform, highly organic and relatively young, the chances of differential preservation conditions along the sequence are low. Additionally, the robust, high-resolution age-depth model that was developed for the Bulusan sequence (Fig. S1) suggests stable sedimentation accumulation over the last 1,400 years. We therefore conclude that taphonomic factors have not played a major role in shaping PARs in the Bulusan sequence.

**Notes S5** Identifying significant periods of change with GAMs

The GAM was used to identify periods of significant environmental change (Simpson 2018). In the GAM, the slope of the trend is potentially different at every point in the time series.

Thus, the GAM answers the question of where in the series the response variable is changing, if at all, by determining whether the first derivative at any time point of the fitted trend is inconsistent with a null hypothesis of no change, given the uncertainty in the estimate of the derivative. The first derivative of the fitted trend is evaluated using finite differences at many points in the time series. Periods of significant change are identified as those time points where the simultaneous confidence interval on the first derivative does not include zero. The intervals strongly guard against false positives, providing confidence in the estimation of the periods of significant change, while the remaining variations in the estimated trend need to be considered carefully to avoid over-interpretation.

Fig. S7 shows the estimated first derivative of the fitted trend in the PAR time series for all identified plant taxa. For most of the taxa, the simultaneous interval on the first derivative either borders zero (*Syzygium*, *Weinmannia*) or excludes zero (*Calamus*, *Elaeocarpus*) at a single or multiple points in the PAR record. For the latter records, we have evidence to reject the null hypothesis of no change. In the case of the Dipterocarp PARs, although the estimated trend suggests a number of oscillations, the estimated trend is sufficiently uncertain that the simultaneous interval on the first derivative includes 0 throughout the record. This is also true for the PAR records of many other taxa (*Caryota*, *Celtis*, *Ficus*, *Macaranga*, *Pinanga* and *Syzygium*). This is suggesting that, while pronounced, changes in PARs in these records are not statistically significant.

**Notes S6** Comparison of GAMs with Gamma and Quasi-Poisson distribution

To evaluate the effect of removing zero values from the PAR records in the GAMs with gamma distribution, we also run the GAMs based on the Quasi-Poisson family that supports the presence of zero values in data but requires the conversion of PAR values to integers. The two GAM family approaches yielded very similar results (see Tables S2, S5-S6, and Fig. S8 for a detailed account).

**Table S1** AMS$^{14}$C dates of the Bulusan sedimentary sequence measured at the NERC Radiocarbon Facility, East Kilbride.

| Lab ID | Sample Material | Core Depth | Age BP | Error |
|---|---|---|---|---|
| SUERC-64001 | Terrestrial plant | 59 | 240 | ±35 |
| SUERC-64002 | Terrestrial plant | 69 | 293 | ±37 |
| SUERC-64003 | Terrestrial plant | 74 | 450 | ±37 |
| SUERC-64005 | Terrestrial plant | 82 | 323 | ±37 |
| SUERC-64004 | Terrestrial plant | 84 | 401 | ±37 |
| SUERC-51111 | Terrestrial plant | 126 | 617 | ±38 |
| SUERC-64006 | Terrestrial plant | 163 | 609 | ±37 |
| SUERC-51112 | Terrestrial plant | 204.5 | 1146 | ±35 |
| SUERC-64010 | Terrestrial plant | 272 | 1235 | ±37 |
| SUERC-64011 | Terrestrial plant | 342.5 | 1629 | ±37 |

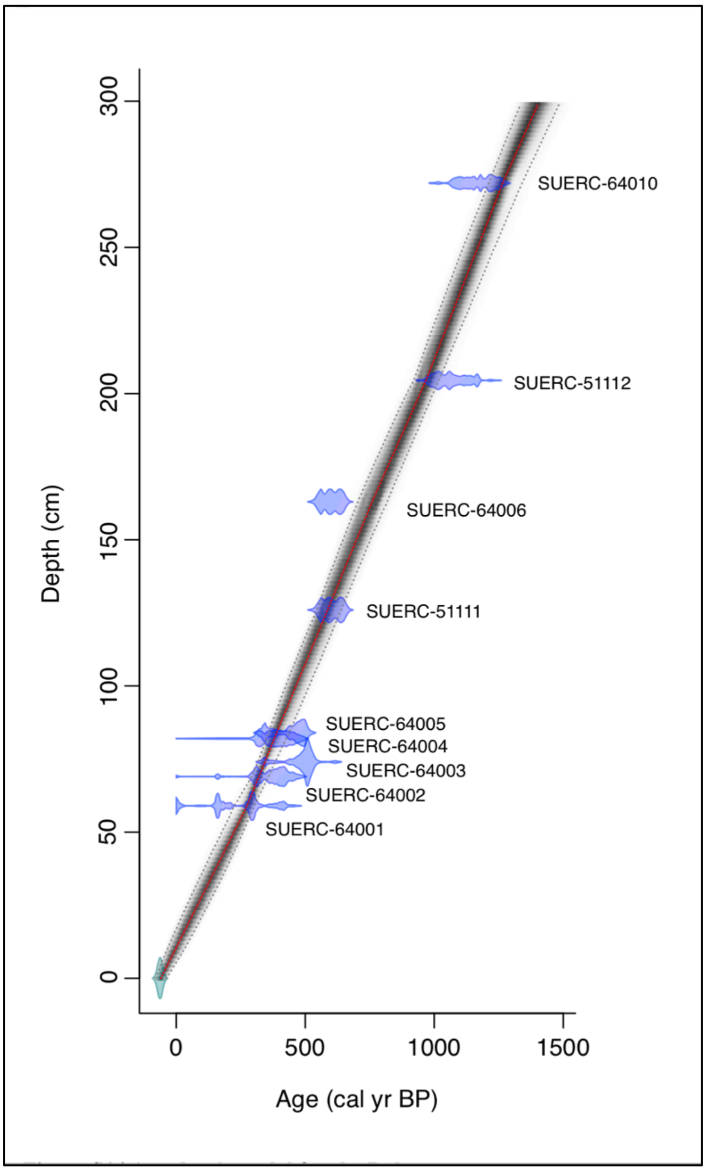


**Fig. S1** Age-depth model for the Bulusan sedimentary sequence (Prohaska *et al.* 2022)**.** The model was constructed in R version 3.2.4 (R Core Team 2020) using *Bacon 2.2* (Blaauw & Christen 2011, 2015) and the IntCal13 radiocarbon calibration curve for the Northern Hemisphere (Reimer *et al.* 2013). Probability distributions for the nine radiocarbon calibrations are shown in blue. The top of the sediment sequence is assigned the year the cores were collected. The shaded areas represent the 1σ probability age ranges by interpolating between $^{14}C$ age control points. The red dashed line represents the age model used in further analysis. The SUERC-64011 date is not shown here as it lies below the analysed section.

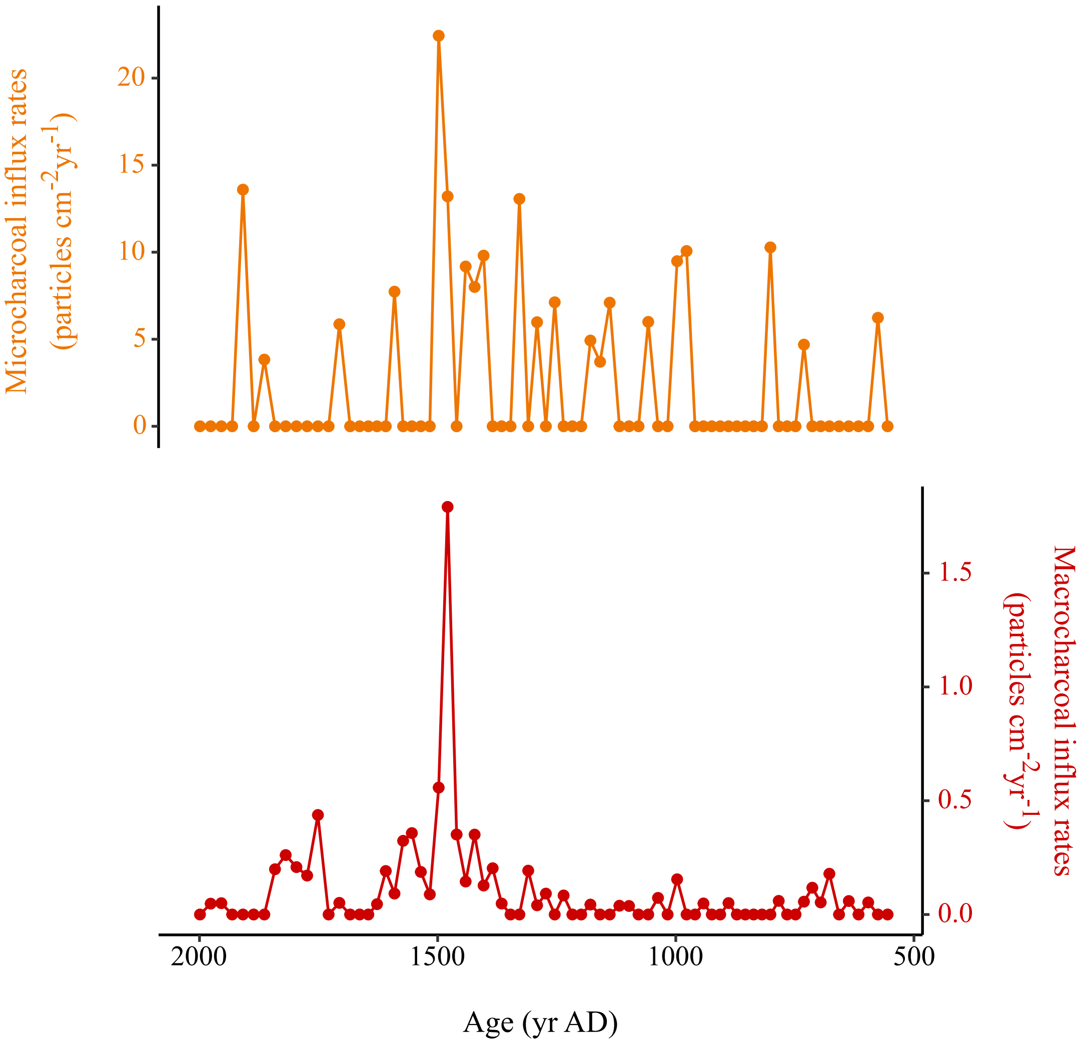


**Fig. S2** Microcharcoal influx rates in relation to macrocharcoal influx rates in the Bulusan record. Both micro- and macro- charcoal influx rates are relatively low throughout the sequence. The peak in macrocharcoal influx rates around ~AD 1500 is not accompanied by an equivalent increase in microcharcoal influx rates, suggesting that this is a local fire signal. Data were plotted with *ggplot2* package (Wickham 2009) in R version 3.2.4 (R Core Team 2020).

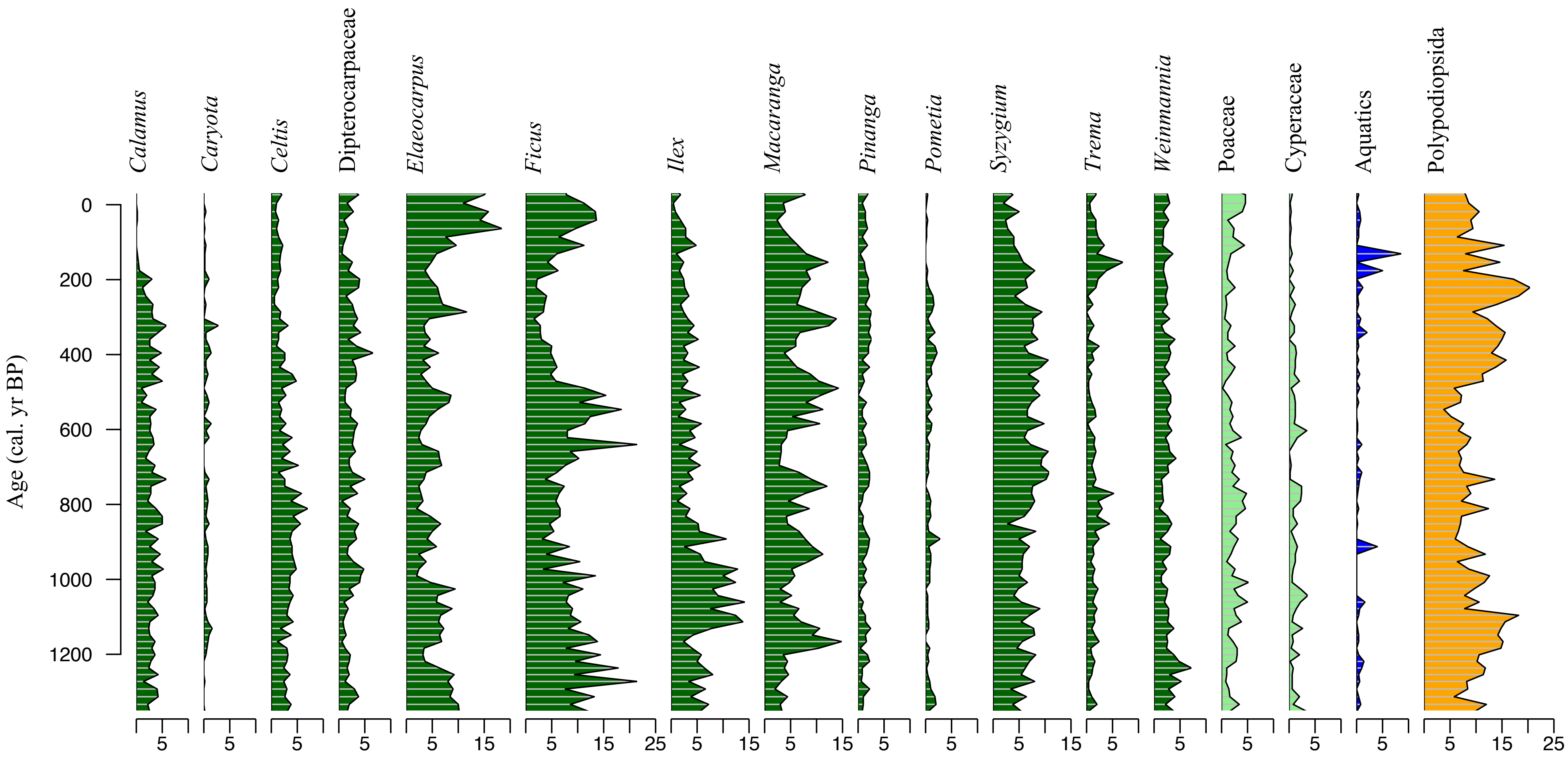


**Fig. S3** Pollen diagram of the sedimentary sequence from Bulusan Lake. Changes in the abundance of the plant taxa identified in the Bulusan record during the last 1,400 years, expressed as pollen percentages. Data was plotted with *rioja (Juggins* 2015*)* in R version 3.2.4 (R Core Team 2020).

**Table S2** Summary of results of the best fitted generalised additive models with a Quasi-Poisson distribution for describing effects of abiotic factors on plant taxa abundance. General model formula: PAR ~ s($\delta D_{wax}$) + s(sqrt(macrochar)) + s(sed$\delta^{15}N$) + s(sed$P_{conc}$), distribution family = Quasi-Poisson, smoothing parameter estimation method = Restricted Maximum Likelihood. PARs = pollen accumulation rates (grains $cm^{-2}yr^{-1}$), macrochar = macrocharcoal influx rates (particles $m^{-2}yr^{-1}$), $\delta D_{wax}$ = $\delta D$ values of the $nC_{29}$ alkane (‰), sed$\delta^{15}N$ = $\delta^{15}N$ values of bulk sediment (‰), sed$P_{conc}$ = phosphorus concentration of bulk sediment (XRF counts per second), POS = significant positive relationship, NEG = significant negative relationship, VAR = significant varying relationship, • = no significant relationship. The fit is considered significant at p-value <0.05. Data was analysed using the mgcv 1.8-31 package (Wood 2018).

| **Plant taxa abundance/ vegetation composition** | **Abiotic factors** | | | |
|---|---|---|---|---|
| | **$\delta D_{wax}$** | **sqrt(macrochar)** | **sed$\delta^{15}N$** | **sed$P_{conc}$** |
| *Calamus* PARs | VAR | • | • | POS |
| *Caryota* PARs | • | • | • | • |
| *Celtis* PARs | • | • | • | • |
| Dipterocarpaceae PARs | NEG | • | • | POS |
| *Elaeocarpus* PARs | NEG | • | • | • |
| *Ficus* PARs | • | VAR | VAR | • |
| *Ilex* PARs | • | • | • | • |
| *Macaranga* PARs | • | POS | • | • |
| *Pinanga* PARs | NEG | • | • | POS |
| *Poaceae* PARs | VAR | • | • | • |
| Polypodiopsida PARs | • | • | • | • |
| *Pometia* PARs | • | • | NEG | • |
| *Syzygium* PARs | • | POS | • | • |
| *Trema* PARs | • | • | • | • |
| *Weinmannia* PARs | • | POS | NEG | • |

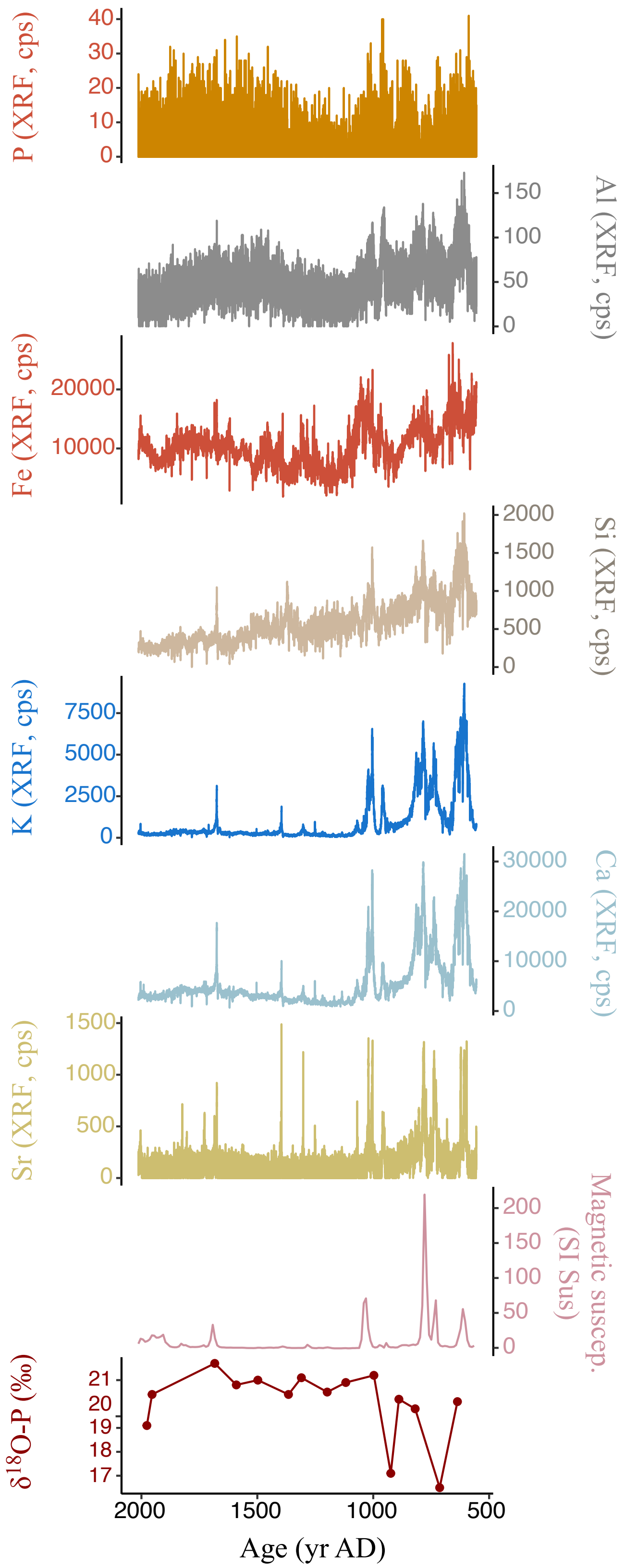


**Fig. S4** Comparison of the record of bulk sediment phosphorus concentrations ($sedP_{conc}$, proxy for phosphorus availability, cps) with the records of volcanic trace elements concentrations (cps), magnetic susceptibility values (SI Sus) and bulk sediment phosphate oxygen isotope values ($\delta^{18}O$-P, ‰) from the Bulusan sedimentary sequence. Data were plotted with *ggplot2* package (Wickham 2009) in R version 3.6.3 (R Core Team 2020).

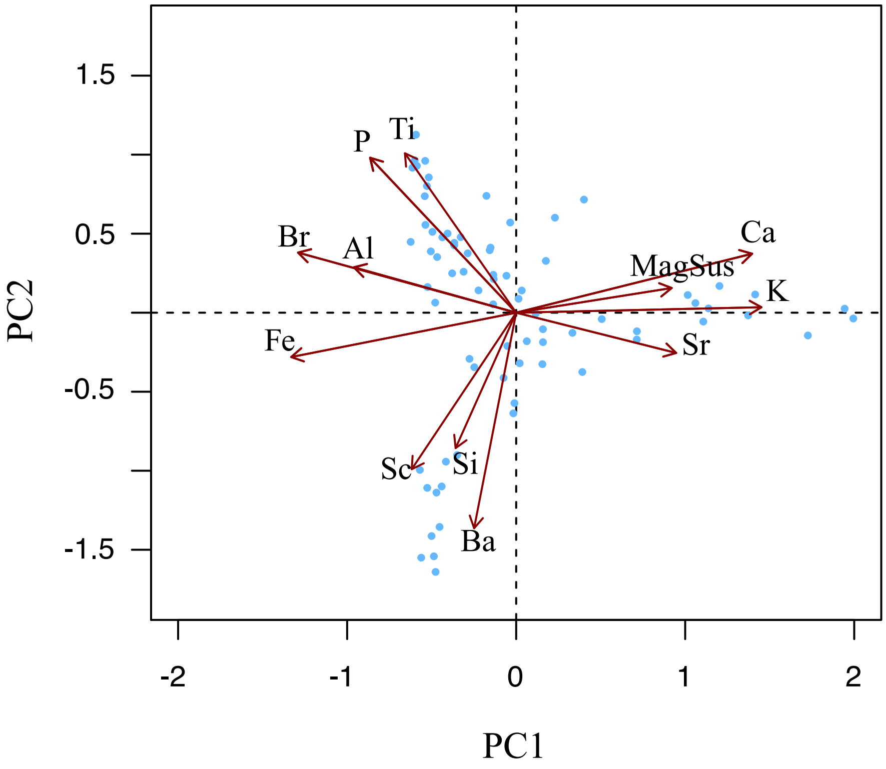


**Fig. S5** PCA results of the abundance dynamics of main geochemical elements measured with the Itrax Core scanner and the magnitude of magnetic susceptibility (MagSus) measured by Barrington MS2 System in the Bulusan sedimentary sequence. Phosphorus (P), potassium (K), calcium (Ca), strontium (Sr), iron (Fe), aluminium (Al), silicon (Si), titanium (Ti), bromine (Br), barium (Ba), and scandium (Sc). The 1$^{st}$ and 2$^{nd}$ axes of the ordination explains 39.9% and 19.8%, respectively, of the variance in the abundance of the elements and the magnitude of magnetic susceptibility over time. Elements Ca, K, Sr, and magnetic susceptibility are clustered together and separated from the other elements by the 1$^{st}$ axis, which suggest their similar dynamics and a common source. Data was plotted with *vegan* package version 2.3-5. (Oksanen *et al.* 2016) in R version 3.2.4 (R Core Team 2020).

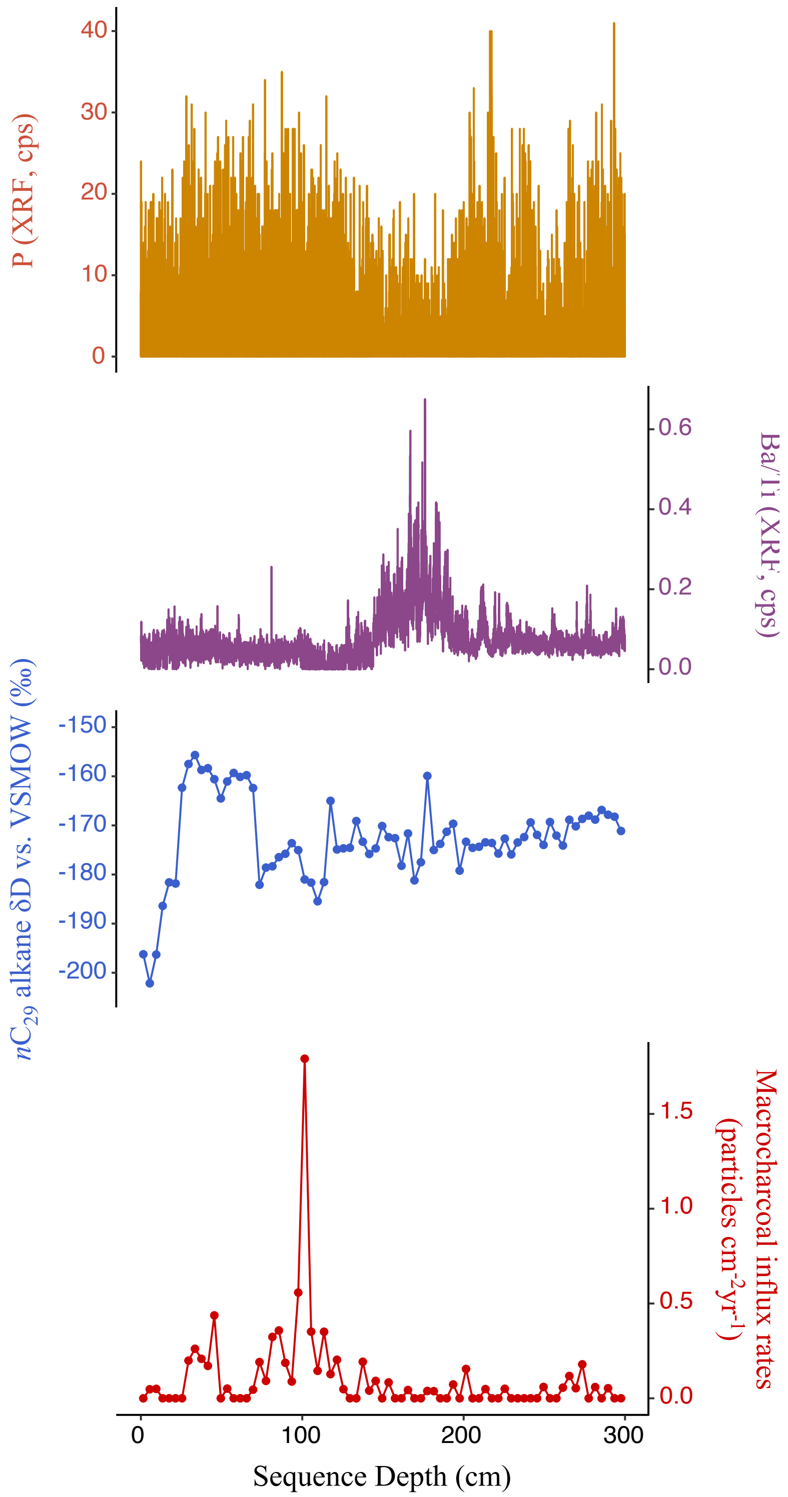


**Fig. S6** Comparison of the record of bulk sediment phosphorus concentrations (sed$P_{conc}$, proxy for phosphorus availability) with the record of $nC_{29}$ alkane δD values (proxy for precipitation amount), macrocharcoal influx rates (proxy for local fire activity), and Ba/Ti ratio (proxy for organic to inorganic matter ratio) from the Bulusan sedimentary sequence. Data were plotted with *ggplot2* package (Wickham 2009) in R version 3.2.4 (R Core Team 2020).

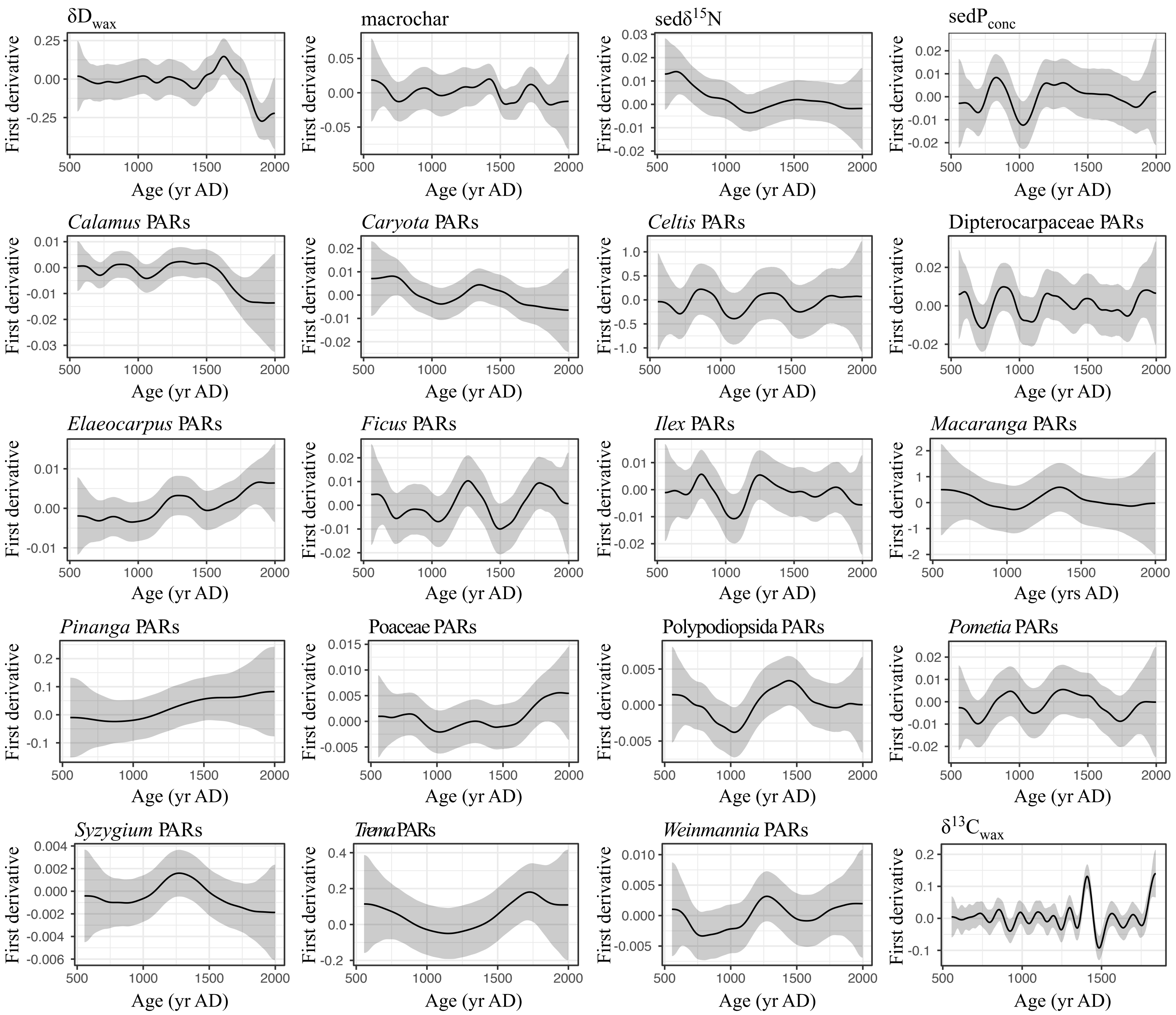

$\delta D_{wax}$
First derivative
0.25
0.00
-0.25
500 1000 1500 2000
Age (yr AD)
macrochar
First derivative
0.05
0.00
-0.05
500 1000 1500 2000
Age (yr AD)
sed$\delta^{15}N$
First derivative
0.03 0.02 0.01 0.00 -0.01 -0.02
500 1000 1500 2000
Age (yr AD)
sed$P_{conc}$
First derivative
0.02 0.01 0.00 -0.01 -0.02
500 1000 1500 2000
Age (yr AD)
*Calamus* PARs
First derivative
0.01 0.00 -0.01 -0.02 -0.03
500 1000 1500 2000
Age (yr AD)
*Caryota* PARs
First derivative
0.02 0.01 0.00 -0.01 -0.02
500 1000 1500 2000
Age (yr AD)
*Celtis* PARs
First derivative
1.0 0.5 0.0 -0.5 -1.0
500 1000 1500 2000
Age (yr AD)
Dipterocarpaceae PARs
First derivative
0.02 0.00 -0.02
500 1000 1500 2000
Age (yr AD)
*Elaeocarpus* PARs
First derivative
0.01 0.00 -0.01
500 1000 1500 2000
Age (yr AD)
*Ficus* PARs
First derivative
0.02 0.01 0.00 -0.01 -0.02
500 1000 1500 2000
Age (yr AD)
*Ilex* PARs
First derivative
0.01 0.00 -0.01 -0.02
500 1000 1500 2000
Age (yr AD)
*Macaranga* PARs
First derivative
2 1 0 -1 -2
500 1000 1500 2000
Age (yrs AD)
*Pinanga* PARs
First derivative
0.2 0.1 0.0 -0.1
500 1000 1500 2000
Age (yr AD)
Poaceae PARs
First derivative
0.015 0.010 0.005 0.000 -0.005
500 1000 1500 2000
Age (yr AD)
Polypodiopsida PARs
First derivative
0.005 0.000 -0.005
500 1000 1500 2000
Age (yr AD)
*Pometia* PARs
First derivative
0.02 0.01 0.00 -0.01 -0.02
500 1000 1500 2000
Age (yr AD)
*Syzygium* PARs
First derivative
0.004 0.002 0.000 -0.002 -0.004 -0.006
500 1000 1500 2000
Age (yr AD)
*Trema* PARs
First derivative
0.4 0.2 0.0 -0.2
500 1000 1500 2000
Age (yr AD)
*Weinmannia* PARs
First derivative
0.010 0.005 0.000 -0.005
500 1000 1500 2000
Age (yr AD)
$\delta^{13}C_{wax}$
First derivative
0.2 0.1 0.0 -0.1
500 1000 1500
Age (yr AD)

**Fig. S7** Significant periods of change in the key paleo-records from Bulusan Lake. Estimated first derivatives (black lines) and 95% simultaneous confidence intervals (grey bands) of the generalised additive model trends with GCV-based smoothness selection (gam = ~ Age) fitted to the individual paleo-records. The models detect significant temporal change in the response variables where the simultaneous interval does not include 0 (Simpson 2018). Models for *Celtis*, *Macaranga*, *Pinanga* and *Trema* were corrected for interval heteroscedacity (sampling intervals). PARs (grains $cm^{-2}yr^{-1}$), macrocharcoal influx rates (particles $m^{-2}yr^{-1}$), $\delta D_{wax}$ = δD values of the $nC_{29}$ alkane (‰), $\delta^{13}C_{wax}$ = $\delta^{13}C$ values of the $nC_{29}$ alkane (‰), sed$\delta^{15}N$ = $\delta^{15}N$ values of bulk sediment (‰), and $sedP_{conc}$ = phosphorus concentrations of bulk sediment (XRF counts per second). Data was analysed using the mgcv 1.8-31 package (Wood 2018) and plotted in R version 3.6.3 (R Core Team 2020).

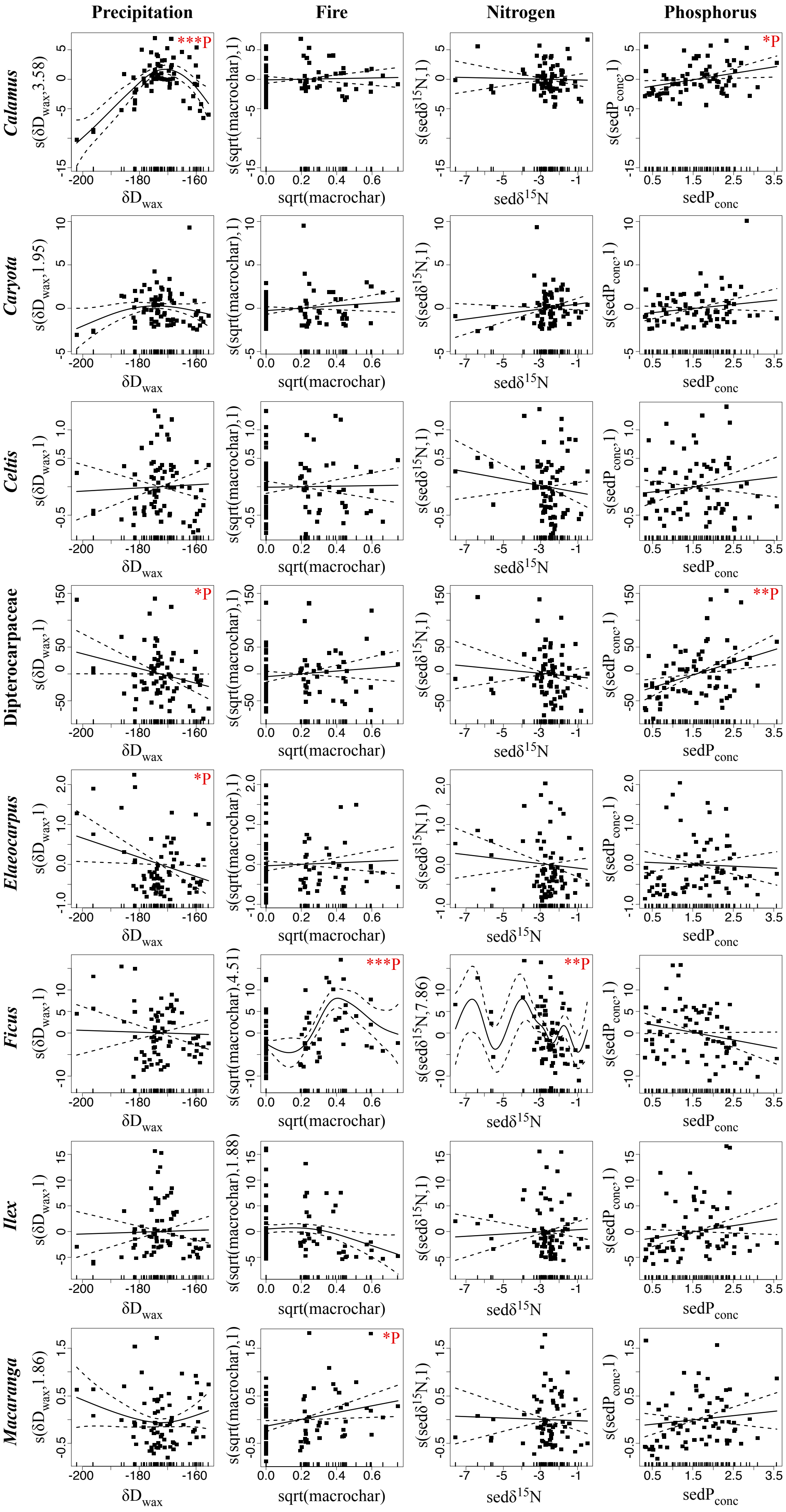

Precipitation
Fire
Nitrogen
Phosphorus
Calamus
Caryota
Celtis
Dipterocarpaceae
Elaeocarpus
Ficus
Ilex
Macaranga
***P
*P
**P
δD_wax
sqrt(macrochar)
sedδ15N
sedP_conc

**Fig. S8** Modelled effects of abiotic factors on plant taxa abundance at Bulusan using generalised additive models with a Quasi-Poisson distribution. Plots of GAM smooth functions from the best fitted models for describing the changes in individual taxa's pollen accumulation rates (proxy for taxa abundance). General model formula: PAR ~ s($\delta D_{wax}$) + s(sqrt(macrochar)) + s(sed$\delta^{15}N$) + s(sed$P_{conc}$), distribution family = Quasi-Poisson, smoothing parameter estimation method = Restricted Maximum Likelihood. PARs = pollen accumulation rates (grains $cm^{-2}yr^{-1}$), macrochar = macrocharcoal influx rates (particles $m^{-2}yr^{-1}$), $\delta D_{wax}$ = $\delta D$ values of the *n*$C_{29}$ alkane (‰), sed$\delta^{15}N$ = $\delta^{15}N$ values of bulk sediment (‰), sed$P_{conc}$ = phosphorus concentration of bulk sediment (XRF counts per second). The tick marks on the x-axis are observed data points. The y-axis represents the partial effect of each variable. The dotted lines indicate the 95% confidence intervals. Asterisks denote significance levels: *P-value $\leq$ 0.05; **P-value $\leq$ 0.01; ***P-value $\leq$ 0.001. Data was analysed and plotted using the mgcv 1.8-31 package (Wood 2018) in R version 3.6.3 (R Core Team 2020).

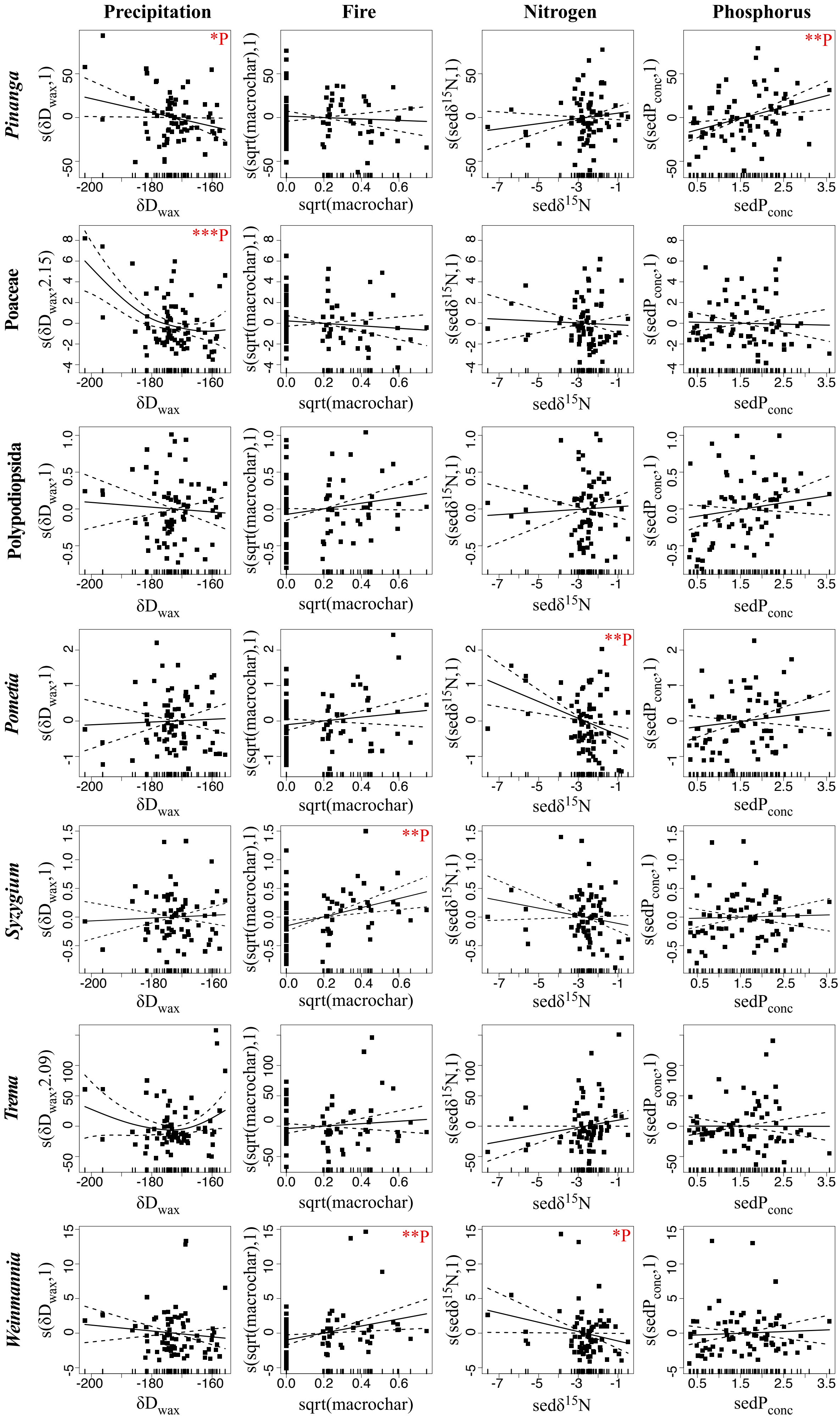

Precipitation
Fire
Nitrogen
Phosphorus
Pinanga
Poaceae
Polypodiopsida
Pometia
Syzygium
Trema
Weinmannia
*P
**P
***P
*P
**P
**P
**P
*P
δD_wax
sqrt(macrochar)
sedδ15N
sedP_conc
s(δD_wax,1)
s(δD_wax,2.15)
s(δD_wax,2.09)
s(sqrt(macrochar),1)
s(sedδ15N,1)
s(sedP_conc,1)

**Fig. S8 (continued)** Modelled effects of abiotic factors on plant taxa abundance at Bulusan using generalised additive models with a Quasi-Poisson distribution.

| *Model* | *Model Structure* | *Distribution family* | *Coefficients (estimate/ SE/ t value/ Pr(>\|t\|)) Intercept* | *Approx. signif. of smooth terms (edf/ Ref. df/ F/ p-value)* | *R-sq.(adj)* | *Scale est.* | -REML | Deviance explained |
|---|---|---|---|---|---|---|---|---|
| $\delta D_{wax}$ | $\delta D_{wax}$ ~ s(Age, k = 30) | gaussian | -172.77/ 0.50/ -343.4/ <2e-16 | 11.92/ 14.76/ 14.68/ <2e-16 | 0.75 | 18.98 | 232.59 | 78.80% |
| macrochar | macrochar ~ s(Age, k = 20) | est. tweedie | -2.90/ 0.16/ -18.3/ <2e-16 | 10.42/ 12.96/ 7.98/ 1.14e-10 | 0.43 | 0.12 | 8.39 | 61.10% |
| sed$\delta^{15}N$ | sed$\delta^{15}N$ ~ s(Age, k = 30) | est. tweedie | -2.73/ 0.11/ -24.98/ <2e-16 | 5.35/ 6.68/ 10.41/ 3.85e-09 | 0.50 | 0.82 | 103.12 | 53.40% |
| sed$P_{conc}$ | sed$P_{conc}$ ~ s(Age, k = 25) | est. tweedie | 0.37/ 0.05/ 8.07/ 2.84e-11 | 11.41/ 14.05/ 8.63/ 2.72e-12 | 0.40 | 0.15 | 78.41 | 63.80% |
| *Calamus* | PARs ~ s(Age, k = 25) | est. tweedie | 4.28/ 0.08/ 51.65/ <2e-16 | 7.76/ 9.68/ 5.65/ 4.48e-06 | 0.44 | 13.94 | 374.22 | 60.80% |
| *Caryota* | PARs ~ s(Age, k = 20) | est. tweedie | 2.57/ 0.11/ 23.59/ <2e-16 | 5.69/ 7.12/ 3.32/ 0.00397 | 0.21 | 8.40 | 243.72 | 28.80% |
| *Celtis* | PARs ~ s(Age, k = 45) | est. tweedie | 4.53/ 0.05/ 96.84/ <2e-16 | 7.70/ 9.62/ 5.21/ 1.09e-05 | 0.41 | 3.50 | 388.24 | 46.20% |
| Dipterocarpaceae | PARs ~ s(Age, k = 40) | est. tweedie | 4.39/ 0.05/ 86.57/ <2e-16 | 12.76/ 15.87/ 3.82/ 4.86e-05 | 0.36 | 0.27 | 388.78 | 52.20% |
| *Elaeocarpus* | PARs ~ s(Age, k = 30) | est. tweedie | 5.39/ 0.06/ 89.55/ <2e-16 | 6.09/ 7.60/ 11.82/ 2.34e-11 | 0.65 | 0.40 | 465.54 | 59.60% |
| *Ficus* | PARs ~ s(Age, k = 30) | est. tweedie | 5.67/ 0.06/ 96.36/ <2e-16 | 10.15/ 12.61/ 8.72/ 1.62e-11 | 0.38 | 0.27 | 492.04 | 61.90% |
| *Ilex* | PARs ~ s(Age, k = 30) | est. tweedie | 4.96/ 0.05/ 92.3/ <2e-16 | 9.65/ 12/ 9.82/ 7.98e-13 | 0.56 | 0.27 | 432.28 | 65.70% |
| *Macaranga* | PARs ~ s(Age, k = 25) | est. tweedie | 5.47/ 0.06/ 94.33/ <2e-16 | 6.43/ 8.02/ 3.66/ 0.00127 | 0.19 | 0.26 | 464.82 | 32.80% |
| *Pinanga* | PARs ~ s(Age, k = 10) | est. tweedie | 3.89/ 0.06/ 65.03/ <2e-16 | 2.60/ 3.24/ 5.71/ 0.00114 | 0.22 | 4.84 | 348.09 | 20.50% |
| Poaceae | PARs ~ s(Age, k = 40) | est. tweedie | 4.29/ 0.05/ 82.7/ <2e-16 | 5.62/ 7.02/ 4.57/ 0.000276 | 0.34 | 0.20 | 371.33 | 36.80% |
| Polypodiopsida | PARs ~ s(Age, k = 30) | est. tweedie | 5.89/ 0.04/ 167.8/ <2e-16 | 6.79/ 8.48/ 9.66/ 6.12e-10 | 0.49 | 3.83 | 468.85 | 56.90% |
| *Pometia* | PARs ~ s(Age, k = 30) | est. tweedie | 3.13/ 0.08/ 37.26/ <2e-16 | 8.18/ 10.20/ 5.24/ 7.11e-06 | 0.40 | 6.67 | 303.87 | 44.50% |
| *Szygium* | PARs ~ s(Age, k = 20) | est. tweedie | 5.46/ 0.05/ 111.9/ <2e-16 | 4.22/ 5.27/ 2.67/ 0.0255 | 0.14 | 0.77 | 454.90 | 19.40% |
| *Trema* | PARs ~ s(Age, k = 20) | est. tweedie | 3.92/ 0.07/ 55.45/ <2e-16 | 4.25/ 5.312/ 4.41/ 0.0013 | 0.22 | 3.05 | 360.00 | 26.30% |
| *Weinmannia* | PARs ~ s(Age, k = 20) | est. tweedie | 4.60/ 0.06/ 71.92/ <2e-16 | 5.58/ 6.96/ 4.48/ 0.000358 | 0.22 | 0.31 | 408.72 | 34.10% |
| $\delta^{13}C_{wax}$ | $\delta^{13}C_{wax}$ ~ s(Age, k = 30) | gaussian | -33.83/ 0.06/ -536/ <2e-16 | 24.77/ 27.58/ 19.73/ <2e-16 | 0.90 | 0.26 | 100.87 | 93.50% |

**Table S3** Description of the best fitted generalised additive models of temporal dynamics of paleo-records from Bulusan Lake with GCV-based smoothness selection. PARs = pollen accumulation rates (grains $cm^{-2}yr^{-1}$), macrochar = macrocharcoal influx rates (particles $m^{-2}yr^{-1}$), $\delta^{13}C_{wax}$ = $\delta^{13}C$ values of the $nC_{29}$ alkane (‰), $\delta DC_{29}$ = δD values of the $nC_{29}$ alkane (‰), sed$\delta^{15}N$ = $\delta^{15}N$ values of bulk sediment (‰), sed$P_{conc}$ = phosphorus concentration of bulk sediment (XRF counts per second), est. tweedie = tweedie family where p is estimated during fitting, Age = calibrated years before the present, REML = Restricted Maximum Likelihood. Data was analysed using the mgcv 1.8-31 package (Wood 2018).

| *Model* | *Model Structure* | *Correlation structure* | *Distribution family* | *Coefficients (estimate/ SE/ t value/ Pr(>\|t\|)) Intercept* | *Approx. signif. of smooth terms (edf/ Ref. df/ F/ p-value)* | *R-sq.(adj)* | *Scale est.* |
|---|---|---|---|---|---|---|---|
| $\delta D_{wax}$ | $\delta D_{wax}$ ~ s(Age, k = 30) | corCAR1(~ Age) | gaussian | -172.77/ 0.50/ -343.4/ <2e-16 | 11.92/ 14.76/ 14.68/ <2e-16 | 0.75 | 18.98 |
| macrochar | macrochar ~ s(Age, k = 26) | corCAR1(~ Age) | fix. tweedie | -2.85/ 0.17/ -17.3/ <2e-16 | 8.5/ 8.5/ 8.75/ 6.79e-09 | 0.38 | 0.20 |
| sed$\delta^{15}N$ | sed$\delta^{15}N$ ~ s(Age, k = 30) | corCAR1(~ Age) | fix. tweedie | -2.73/ 0.10/ -27.96/ <2e-16 | 5.54/ 5.54/ 11.14/ 1.31e-08 | 0.47 | 0.68 |
| sed$P_{conc}$ | sed$P_{conc}$ ~ s(Age, k = 25) | corCAR1(~ Age) | fix. tweedie | 0.40/ 0.05/ 8.00/ 3.13e-11 | 9.51/ 9.51/ 6.78/ 2.8e-07 | 0.41 | 0.25 |
| *Calamus* | PARs ~ s(Age, k = 27) | corCAR1(~ Age) | fix. tweedie | 4.27/ 0.07/ 61.87/ <2e-16 | 7.43/ 7.43/ 10.31/ 1.02e-09 | 0.49 | 5.56 |
| *Caryota* | PARs ~ s(Age, k = 21) | corCAR1(~ Age) | fix. tweedie | 2.58/ 0.11/ 23.01/ <2e-16 | 4.94/ 4.94/ 4.23/ 0.00293 | 0.17 | 6.10 |
| *Celtis* | PARs ~ s(Age, k = 39) | corCAR1(~ Age) | fix. tweedie | 4.54/ 0.05/ 89.27/ <2e-16 | 1/ 1/ 28.55/ 8.4e-07 | 0.28 | 5.67 |
| Dipterocarpaceae | PARs ~ s(Age, k = 68) | corCAR1(~ Age) | fix. tweedie | 4.42/ 0.05/ 85.8/ <2e-16 | 11.53/ 11.53/ 3.64/ 0.000376 | 0.32 | 5.32 |
| *Elaeocarpus* | PARs ~ s(Age, k = 44) | corCAR1(~ Age) | fix. tweedie | 5.42/ 0.06/ 84.39/ <2e-16 | 4.80/ 4.80/ 22.61/ 2.05e-15 | 0.64 | 16.58 |
| *Ficus* | PARs ~ s(Age, k = 30) | corCAR1(~ Age) | fix. tweedie | 5.70/ 0.07/ 84/ <2e-16 | 9.49/ 9.49/ 7.82/ 1.93e-08 | 0.39 | 22.76 |
| *Ilex* | PARs ~ s(Age, k = 30) | corCAR1(~ Age) | fix. tweedie | 4.98/ 0.06/ 87.2/ <2e-16 | 8.86/ 8.86/ 13.29/ 3.9e-14 | 0.57 | 9.43 |
| *Macaranga* | PARs ~ s(Age, k = 39) | corCAR1(~ Age) | fix. tweedie | 5.51/ 0.07/ 81.6/ <2e-16 | 1/ 1/ 6.68/ 0.0117 | 0.07 | 20.88 |
| *Pinanga* | PARs ~ s(Age, k = 10) | corCAR1(~ Age) | fix. tweedie | 3.89/ 0.06/ 68.03/ <2e-16 | 2.30/ 2.30/ 8.24/ 0.000325 | 0.22 | 4.45 |
| Poaceae | PARs ~ s(Age, k = 40) | corCAR1(~ Age) | fix. tweedie | 4.30/ 0.05 80.3/ <2e-16 | 5.05/ 5.05/ 9.27/ 4.68e-07 | 0.37 | 5.24 |
| Polypodiopsida | PARs ~ s(Age, k = 27) | corCAR1(~ Age) | fix. tweedie | 5.89/ 0.03/ 172/ <2e-16 | 6.66/ 6.66/ 12.44/ 7.5e-11 | 0.49 | 7.01 |
| *Pometia* | PARs ~ s(Age, k = 30) | corCAR1(~ Age) | fix. tweedie | 3.13/ 0.08/ 38.51/ <2e-16 | 7.87/ 7.87/ 6.50/ 2.36e-06 | 0.39 | 4.79 |
| *Szygium* | PARs ~ s(Age, k = 11) | corCAR1(~ Age) | fix. tweedie | 5.47/ 0.05/ 109.1/ <2e-16 | 3.68/ 3.68/ 3.15/ 0.0399 | 0.13 | 11.19 |
| *Trema* | PARs ~ s(Age, k = 20) | corCAR1(~ Age) | fix. tweedie | 3.92/ 0.07/ 52.55/ <2e-16 | 3.49/ 3.49/ 6.34/ 0.000784 | 0.21 | 7.70 |
| *Weinmannia* | PARs ~ s(Age, k = 13) | corCAR1(~ Age) | fix. tweedie | 4.63/ 0.07/ 66.83/ <2e-16 | 2.87/ 2.87/ 5.85/ 0.00318 | 0.15 | 11.36 |
| $\delta^{13}C_{wax}$ | $\delta^{13}C_{wax}$ ~ s(Age, k = 30) | corCAR1(~ Age) | gaussian | -33.83/ 0.06/ -536/ <2e-16 | 4.77/ 24.77/ 21.64/ <2e-16 | 0.90 | 0.26 |

**Table S4** Description of the best fitted generalised additive models of temporal dynamics of paleo-records from Bulusan Lake with a continuous-time AR(1) process estimated using REML smoothness selection. PARs = pollen accumulation rates (grains $cm^{-2}yr^{-1}$), macrochar = macrocharcoal influx rates (particles $m^{-2}yr^{-1}$), $\delta^{13}C_{wax}$ = $\delta^{13}C$ values of the $nC_{29}$ alkane (‰), $\delta D_{wax}$ = $\delta D$ values of the $nC_{29}$ alkane (‰), sed$\delta^{15}N$ = $\delta^{15}N$ values of bulk sediment (‰), sed$P_{conc}$ = phosphorus concentration of bulk sediment (XRF counts per second), fix.tweedie = tweedie family where p is fixed, Age = calibrated years before the present, REML = Restricted Maximum Likelihood. Data was analysed using the mgcv 1.8-31 package (Wood 2018).

| *Model* | *Model Structure* | *Correlation structure* | *Distribution family (link)* | *Approx. signif. of smooth terms (edf/Ref.df/F/p-value)* | | | | *R-sq.(adj)* | *Scale est.* |
|---|---|---|---|---|---|---|---|---|---|
| | | | | **$\delta D_{wax}$** | **sqrt(macrochar)** | **$sed\delta^{15}N$** | **$sedP_{conc}$** | | |
| *Calamus*^ | $PAR \sim s(\delta D_{wax}) + s(sqrt(macrochar)) + s(sed\delta^{15}N) + s(sedP_{conc})$ | / | Gamma (log) | 3.21/ 3.21/ 7.17/ 0.0003 *** | 1.00/ 1.00/ 0.01/ 0.9279 | 1.00/ 1.00/ 0.16/ 0.6923 | 1.00/ 1.00/ 6.27/ 0.0150 * | 0.18 | 0.24 |
| *Caryota*^ | $PAR \sim s(\delta D_{wax}) + s(sqrt(macrochar)) + s(sed\delta^{15}N) + s(sedP_{conc})$ | / | Gamma (log) | 1.00/ 1.00/ 1.72/ 0.1962 | 1.00/ 1.00/ 1.35/ 0.2518 | 1.00/ 1.00/ 0.01/ 0.9348 | 1.00/ 1.00/ 5.18/ 0.0274 * | 0.07 | 0.40 |
| *Celtis* | $PAR \sim s(\delta D_{wax}) + s(sqrt(macrochar)) + s(sed\delta^{15}N) + s(sedP_{conc})$ | / | Gamma (log) | 2.30/ 2.30/ 2.02/ 0.2470 | 1.00/ 1.00/ 0.12/ 0.7330 | 1.00/ 1.00/ 1.19/ 0.280 | 1.00/ 1.00/ 0.05/ 0.8170 | 0.05 | 0.24 |
| Dipterocarpaceae | $PAR \sim s(\delta D_{wax}) + s(sqrt(macrochar)) + s(sed\delta^{15}N) + s(sedP_{conc})$ | corExp(~Age) | Gamma (log) | 1.00/ 1.00/ 4.35/ 0.0407 * | 1.00/ 1.00/ 0.91/ 0.3434 | 1.00/ 1.00/ 1.05/ 0.3090 | 1.00/ 1.00/ 10.25/ 0.0021 ** | 0.19 | 0.23 |
| *Elaeocarpus* | $PAR \sim s(\delta D_{wax}) + s(sqrt(macrochar)) + s(sed\delta^{15}N) + s(sedP_{conc})$ | corExp(~Age) | Gamma (log) | 1.00/ 1.00/ 4.52/ 0.0371 * | 1.00/ 1.00/ 1.57/ 0.2140 | 1.00/ 1.00/ 0.32/ 0.5722 | 1.81/ 1.81/ 2.03/ 0.2146 | 0.14 | 0.51 |
| *Ficus* | $PAR \sim s(\delta D_{wax}) + s(sqrt(macrochar)) + s(sed\delta^{15}N) + s(sedP_{conc})$ | corExp(~Age) | Gamma (log) | 1.00/ 1.00/ 3.89/ 0.0528 . | 3.68/ 3.68/ 7.04/ 8.99e-05 *** | 1.00/ 1.00/ 2.81/ 0.0986 . | 1.00/ 1.00/ 1.41/ 0.2395 | 0.23 | 0.39 |
| *Ilex* | $PAR \sim s(\delta D_{wax}) + s(sqrt(macrochar)) + s(sed\delta^{15}N) + s(sedP_{conc})$ | corExp(~Age) | Gamma (log) | 1.00/ 1.00/ 0.26/ 0.6126 | 2.07/ 2.07/ 4.93/ 0.0103 * | 1.00/ 1.00/ 0.23/ 0.6298 | 1.00/ 1.00/ 2.17/ 0.1456 | -0.01 | 0.54 |
| *Macaranga* | $PAR \sim s(\delta D_{wax}) + s(sqrt(macrochar)) + s(sed\delta^{15}N) + s(sedP_{conc})$ | corExp(~Age) | Gamma (log) | 1.00/ 1.00/ 0.48/ 0.4901 | 1.00/ 1.00/ 7.10/ 0.0096 ** | 1.00/ 1.00/ 0.11/ 0.7427 | 1.00/ 1.00/ 2.81/ 0.0984 . | 0.11 | 0.27 |
| *Pinanga* ^‡ | $PAR \sim s(\delta D_{wax}) + s(sqrt(macrochar)) + s(sed\delta^{15}N) + s(sedP_{conc})$ | / | Gamma (log) | 1.00/ 1.00/ 6.25/ 0.0149 * | 1.00/ 1.00/ 0.12/ 0.7296 | 1.00/ 1.00/ 1.93/ 0.1694 | 1.00/ 1.00/ 10.94/ 0.0015 ** | 0.16 | 0.22 |
| Poaceae‡ | $PAR \sim s(\delta D_{wax}) + s(sqrt(macrochar)) + s(sed\delta^{15}N) + s(sedP_{conc})$ | / | Gamma (log) | 8.27/ 8.27/ 4.15/ 0.0004 *** | 1.00/ 1.00/ 0.69/ 0.4109 | 1.00/ 1.00/ 0.13/ 0.7209 | 1.00/ 1.00/ 0.24/ 0.6289 | 0.34 | 0.26 |
| Polypodiopsida | $PAR \sim s(\delta D_{wax}) + s(sqrt(macrochar)) + s(sed\delta^{15}N) + s(sedP_{conc})$ | corExp(~Age) | Gamma (log) | 1.00/ 1.00/ 0.60/ 0.4407 | 1.00/ 1.00/ 4.30/ 0.0418 * | 1.00/ 1.00/ 0.28/ 0.6012 | 1.00/ 1.00/ 2.36/ 0.1291 | 0.05 | 0.16 |
| *Pometia*^ | $PAR \sim s(\delta D_{wax}) + s(sqrt(macrochar)) + s(sed\delta^{15}N) + s(sedP_{conc})$ | / | Gamma (log) | 1.00/ 1.00/ 0.02/ 0.8985 | 1.00/ 1.00/ 4.27/ 0.0432 * | 2.89/ 2.89/ 3.72/ 0.0323 * | 1.00/ 1.00/ 7.29/ 0.0090 ** | 0.29 | 0.33 |
| *Szygium* | $PAR \sim s(\delta D_{wax}) + s(sqrt(macrochar)) + s(sed\delta^{15}N) + s(sedP_{conc})$ | corExp(~Age) | Gamma (log) | 1.00/ 1.00/ 0.26/ 0.6089 | 1.00/ 1.00/ 10.06/ 0.0023 ** | 1.00/ 1.00/ 2.10/ 0.1517 | 1.00/ 1.00/ 0.35/ 0.5583 | 0.11 | 0.17 |
| *Trema*^ | $PAR \sim s(\delta D_{wax}) + s(sqrt(macrochar)) + s(sed\delta^{15}N) + s(sedP_{conc})$ | / | Gamma (log) | 2.08/ 2.08/ 2.63/ 0.0939 . | 1.00/ 1.00/ 0.43/ 0.5153 | 1.00/ 1.00/ 4.40/ 0.0398 * | 1.00/ 1.00/ 0.00/ 0.9956 | 0.20 | 0.41 |
| *Weinmannia* | $PAR \sim s(\delta D_{wax}) + s(sqrt(macrochar)) + s(sed\delta^{15}N) + s(sedP_{conc})$ | corExp(~Age) | Gamma (log) | 1.00/ 1.00/ 1.32/ 0.2555 | 1.00/ 1.00/ 7.40/ 0.0083 ** | 1.00/ 1.00/ 3.741/ 0.0572 . | 1.00/ 1.00/ 0.57/ 0.4523 | 0.07 | 0.36 |
| $\delta^{13}C_{wax}$ | $\delta^{13}C_{wax} \sim s(\delta D_{wax}) + s(sqrt(macrochar)) + s(sed\delta^{15}N) + s(sedP_{conc})$ | corExp(~Age) | Gaussian (identity) | 3.29/ 3.29/ 6.26/ 0.0008 *** | 1.00/ 1.00/ 2.51/ 0.1186 | 1.35/ 1.35/ 0.10/ 0.8345 | 1.00/ 1.00/ 0.12/ 0.7273 | 0.42 | 1.72 |

**Table S5** Description of the best fitted generalised additive models with a Gamma distribution for the effects of abiotic factors on plant taxa abundance (pollen accumulation rates, PARs) and C3/ C4 vegetation composition ($\delta^{13}C$ values of the $nC_{29}$ alkane, $\delta^{13}C_{wax}$) at Bulusan. PARs = pollen accumulation rates (grains $cm^{-2}yr^{-1}$), macrochar = macrocharcoal influx rates (particles $m^{-2}yr^{-1}$), $\delta^{13}C_{wax}$ = $\delta^{13}C$ values of the $nC_{29}$ alkane (*‰*), $\delta D_{wax}$ = $\delta D$ values of the $nC_{29}$ alkane (*‰*), $sed\delta^{15}N$ = $\delta^{15}N$ values of bulk sediment (*‰*), $sedP_{conc}$ = phosphorus concentration of bulk sediment (XRF counts per second), Age = calibrated years before the present, REML = Restricted Maximum Likelihood. The tick marks on the x-axis are observed data points. The y-axis represents the partial effect of each variable. The dotted lines indicate the 95% confidence intervals. ‡ = model cannot converge. ^ = dataset contains zeros, zeros removed prior to running GAMs. Data was analysed and plotted using the mgcv 1.8-31 package (Wood 2018) in R version 3.6.3 (R Core Team 2020).

| *Model* | *Model Structure* | *Correlation structure* | *Distribution family (link)* | *Approx. signif. of smooth terms (edf/Ref.df/F/p-value)* | | | | *R-sq.(adj)* | *Scale est.* |
|---|---|---|---|---|---|---|---|---|---|
| | | | | **$\delta D_{wax}$** | **sqrt(macrochar)** | **sed$\delta^{15}$N** | **sedP$_{conc}$** | | |
| *Calamus*^ | PAR ~ s($\delta D_{wax}$) + s(sqrt(macrochar)) + s(sed$\delta^{15}$N) + s(sedP$_{conc}$) | / | Quasi-Poisson (sqrt) | 3.58/ 3.58/ 17.77/ 8.31e-10 *** | 1.00/ 1.00/ 0.127/ 0.7226 | 1.00/ 1.00/ 0.06/ 0.8067 | 1.00/ 1.00/ 5.88/ 0.0181 * | 0.32 | 24.90 |
| *Caryota*^ | PAR ~ s($\delta D_{wax}$) + s(sqrt(macrochar)) + s(sed$\delta^{15}$N) + s(sedP$_{conc}$) | / | Quasi-Poisson (sqrt) | 1.95/ 1.95/ 2.68/ 0.0807 . | 1.00/ 1.00/ 1.53/ 0.2203 | 1.00/ 1.00/ 2.08/ 0.1536 | 1.00/ 1.00/ 2.03/ 0.1592 | 0.06 | 14.41 |
| *Celtis* | PAR ~ s($\delta D_{wax}$) + s(sqrt(macrochar)) + s(sed$\delta^{15}$N) + s(sedP$_{conc}$) | corExp(~Age) | Quasi-Poisson (log) | 1.00/ 1.00/ 0.11/ 0.740 | 1.00/ 1.00/ 0.03/ 0.867 | 1.00/ 1.00/ 1.30/ 0.2580 | 1.00/ 1.00/ 0.93/ 0.3370 | -0.05 | 24.51 |
| Dipterocarpaceae | PAR ~ s($\delta D_{wax}$) + s(sqrt(macrochar)) + s(sed$\delta^{15}$N) + s(sedP$_{conc}$) | corExp(~Age) | Quasi-Poisson (identity) | 1.00/ 1.00/ 4.06/ 0.0480 * | 1.00/ 1.00/ 1.02/ 0.3157 | 1.00/ 1.00/ 0.57/ 0.4545 | 1.00/ 1.00/ 10.17/ 0.0022 ** | 0.18 | 20.21 |
| *Elaeocarpus* | PAR ~ s($\delta D_{wax}$) + s(sqrt(macrochar)) + s(sed$\delta^{15}$N) + s(sedP$_{conc}$) | corExp(~Age) | Quasi-Poisson (log) | 1.00/ 1.00/ 5.13/ 0.0267 * | 1.00/ 1.00/ 0.40/ 0.5306 | 1.00/ 1.00/ 0.80/ 0.3733 | 1.00/ 1.00/ 0.22/ 0.6406 | 0.14 | 153.15 |
| *Ficus* | PAR ~ s($\delta D_{wax}$) + s(sqrt(macrochar)) + s(sed$\delta^{15}$N) + s(sedP$_{conc}$) | corExp(~Age) | Quasi-Poisson (sqrt) | 1.00/ 1.00/ 0.06/ 0.8098 | 4.51/ 4.51/ 13.25/ 1.02e-08 *** | 7.86/ 7.86/ 3.50/ 0.0015 ** | 1.00/ 1.00/ 3.67/ 0.0602 . | 0.27 | 138.42 |
| *Ilex* | PAR ~ s($\delta D_{wax}$) + s(sqrt(macrochar)) + s(sed$\delta^{15}$N) + s(sedP$_{conc}$) | corExp(~Age) | Quasi-Poisson (sqrt) | 1.00/ 1.00/ 0.07/ 0.7947 | 1.88/ 1.88/ 4.26/ 0.0511 . | 1.00/ 1.00/ 0.24/ 0.6252 | 1.00/ 1.00/ 2.62/ 0.1100 | 0.00 | 95.63 |
| *Macaranga* | PAR ~ s($\delta D_{wax}$) + s(sqrt(macrochar)) + s(sed$\delta^{15}$N) + s(sedP$_{conc}$) | corExp(~Age) | Quasi-Poisson (log) | 1.86/ 1.86/ 1.82/ 0.1994 | 1.00/ 1.00/ 5.97/ 0.0172 * | 1.00/ 1.00/ 0.07/ 0.7937 | 1.00/ 1.00/ 0.94/ 0.3369 | 0.18 | 68.61 |
| *Pinanga*^ | PAR ~ s($\delta D_{wax}$) + s(sqrt(macrochar)) + s(sed$\delta^{15}$N) + s(sedP$_{conc}$) | / | Quasi-Poisson (identity) | 1.00/ 1.00/ 4.45/ 0.0386 * | 1.00/ 1.00/ 0.28/ 0.6002 | 1.00/ 1.00/ 1.80/ 0.1841 | 1.00/ 1.00/ 9.77/ 0.0026 ** | 0.12 | 12.28 |
| Poaceae | PAR ~ s($\delta D_{wax}$) + s(sqrt(macrochar)) + s(sed$\delta^{15}$N) + s(sedP$_{conc}$) | / | Quasi-Poisson (sqrt) | 2.15/ 2.15/ 8.49/ 0.0003 *** | 1.00/ 1.00/ 0.79/ 0.3768 | 1.00/ 1.00/ 0.15/ 0.7048 | 1.00/ 1.00/ 0.06/ 0.8157 | 0.30 | 20.15 |
| Polypodiopsida | PAR ~ s($\delta D_{wax}$) + s(sqrt(macrochar)) + s(sed$\delta^{15}$N) + s(sedP$_{conc}$) | corExp(~Age) | Quasi-Poisson (log ) | 1.00/ 1.00/ 0.26/ 0.6097 | 1.00/ 1.00/ 3.44/ 0.0679 . | 1.00/ 1.00/ 0.17/ 0.6832 | 1.00/ 1.00/ 1.88/ 0.1753 | 0.06 | 58.09 |
| *Pometia* ‡^ | PAR ~ s($\delta D_{wax}$) + s(sqrt(macrochar)) + s(sed$\delta^{15}$N) + s(sedP$_{conc}$) | corExp(~Age) | Quasi-Poisson (log) | 1.00/ 1.00/ 0.10/ 0.7503 | 1.00/ 1.00/ 1.61/ 0.2091 | 1.00/ 1.00/ 10.96/ 0.0015 ** | 1.00/ 1.00/ 1.27/ 0.2643 | 0.16 | 14.48 |
| *Szygium* | PAR ~ s($\delta D_{wax}$) + s(sqrt(macrochar)) + s(sed$\delta^{15}$N) + s(sedP$_{conc}$) | corExp(~Age) | Quasi-Poisson (log) | 1.00/ 1.00/ 0.18/ 0.6727 | 1.00/ 1.00/ 11.16/ 0.0014 ** | 1.00/ 1.00/ 2.83/ 0.0971 . | 1.00/ 1.00/ 0.07/ 0.7901 | 0.12 | 41.06 |
| *Trema*^ | PAR ~ s($\delta D_{wax}$) + s(sqrt(macrochar)) + s(sed$\delta^{15}$N) + s(sedP$_{conc}$) | / | Quasi-Poisson (identity) | 2.09/ 2.09/ 2.44/ 0.1112 | 1.00/ 1.00/ 0.92/ 0.3422 | 1.00/ 1.00/ 3.96/ 0.0508 . | 1.00/ 1.00/ 0.00/ 0.9823 | 0.15 | 23.18 |
| *Weinmannia* | PAR ~ s($\delta D_{wax}$) + s(sqrt(macrochar)) + s(sed$\delta^{15}$N) + s(sedP$_{conc}$) | / | Quasi-Poisson (sqrt) | 1.00/ 1.00/ 0.91/ 0.3436 | 1.00/ 1.00/ 7.12/ 0.0095 ** | 1.00/ 1.00/ 4.28/ 0.0425 * | 1.00/ 1.00/ 0.21/ 0.6503 | 0.08 | 40.20 |

**Table S6** Description of the best fitted generalised additive models with a Quasi-Poisson distribution for the effects of abiotic factors on plant taxa abundance (pollen accumulation rates, PARs) at Bulusan. PARs = pollen accumulation rates (grains $cm^{-2}yr^{-1}$), macrochar = macrocharcoal influx rates (particles $m^{-2}yr^{-1}$), $\delta D_{wax}$ = δD values of the $n$C$_{29}$ alkane (*‰*), sed$\delta^{15}$N = $\delta^{15}$N values of bulk sediment (*‰*), sedP$_{conc}$ = phosphorus concentration of bulk sediment (XRF counts per second), Age = calibrated years before the present, REML = Restricted Maximum Likelihood. The tick marks on the x-axis are observed data points. The y-axis represents the partial effect of each variable. The dotted lines indicate the 95% confidence intervals. ‡ = model cannot converge. ^ = dataset contains zeros, zeros removed prior to running GAMs. Data was analysed and plotted using the mgcv 1.8-31 package (Wood 2018) in R version 3.6.3 (R Core Team 2020).
